# Evaluating drought tolerance stability in soybean by the response of irrigation change captured from time-series multispectral data

**DOI:** 10.1101/2023.04.05.535659

**Authors:** Kengo Sakurai, Yusuke Toda, Kosuke Hamazaki, Yoshihiro Ohmori, Yuji Yamasaki, Hirokazu Takahashi, Hideki Takanashi, Mai Tsuda, Hisashi Tsujimoto, Akito Kaga, Mikio Nakazono, Toru Fujiwara, Hiroyoshi Iwata

## Abstract

This study investigated a method to evaluate the drought tolerance stability of a genotype in a single environmental trial by capturing the plant response to irrigation changes. Genotypes that exhibit stable phenotypes under various drought stress conditions are required for stable crop production. However, considerable time and money are required to evaluate the environmental stability of a genotype through multiple environmental trials. As an index of drought tolerance stability, we calculated the coefficient of variation (CV) of shoot fresh weight of 178 soybean (*Glycine max* (L.) Merr.) accessions in a total of nine types of drought treatments, including changing irrigation treatments (every five or ten days) over 3-year trials. To capture the plant responses to irrigation changes, time-series multispectral (MS) data were collected, including the timings of the irrigation/non-irrigation switch in the changing irrigation treatments. We built a random regression model (RRM) for each of the nine treatments using the time-series MS data. We built a genomic prediction model (MT_RRM_ model) using the genetic random regression coefficients of RRM as secondary traits and evaluated the accuracy of each model for predicting CV. In two out of the three years, the prediction accuracy of MT_RRM_ models built in the changing irrigation treatment was higher than that in the continuous drought treatment in the same year. When the CV was predicted using the MT_RRM_ model across years in the changing irrigation treatment, the prediction accuracy was 61% higher than that of the simple genomic prediction model. These results suggest that drought tolerance stability can be evaluated in a single environmental trial, which may reduce the time and cost of selecting genotypes with high drought tolerance stability.

## 1. Introduction

Soybean (*Glycine max* (L.) Merr.) exhibits a 40% reduction in yield due to drought [1], and genetic improvement of drought tolerance in soybeans is needed. Plant responses to drought stress are associated with drought tolerance [2,3], and phenotypic data can be collected non-destructively from plants using high-throughput phenotyping (HTP). The relationship between spectral reflectance collected by hyperspectral (HS) and multispectral (MS) cameras and drought stress has been reported in several studies [4–6]. Relationships between the normalized difference vegetation index (NDVI), which is calculated from the reflectance of near-infrared and red spectra, and the level of wilting have been reported in soybeans [7]. In soybeans, genotypes with slow-wilting traits exhibit high yield under drought conditions [2,8]. Among several vegetation indices (VIs), normalized difference red-edge (NDRE), calculated from the reflectance of red-edge and red spectra, is reported to be the best vegetation index for detecting drought stress [9]. These plant responses to irrigation changes are continuous, and time-series data can be collected using HTP [10]. New insights can be obtained by capturing and analyzing time-series plant responses to irrigation changes, which are collected using HTP [11]. Chen et al. [12] collected time-series changes in the digital volume of barley calculated from plant images collected using HTP during drought and recovery periods. The speed of recovery differs among genotypes; the faster the speed of recovery, the larger the final biomass [12].

The random regression model (RRM) has been widely used for genetic analysis using time-series data [13–15]. In the RRM, the covariance between each time point in a multivariate mixed model is modeled using Legendre polynomials and spline functions based on the assumption that time-series data are changing continuously [16]. The RRM makes it possible to describe time-series random genetic effects using a small number of parameters [17,18]. The calculated genetic random regression coefficients are used in a genome-wide association study (GWAS) to search for new genes [19] or as secondary traits in a multi-trait model (MTM) to increase the prediction accuracy of the target trait [20].

In terms of drought tolerance, one of the important breeding targets is the drought tolerance stability, which is a stable phenotype under various drought levels (i.e., severe, moderate, and mild) because the amount of rainfed and water availability varies among sites and years [21–25]. Coefficient of variation (CV) for target traits calculated from multi-environmental trials has long been used as an indicator for environmental stability [26–31]. CV is calculated by dividing the mean value of a target trait by its standard deviation, with smaller values indicating greater stability [32]. However, to evaluate the CV of drought tolerance in a given genotype, cultivation trials must be conducted at various drought levels. It is time-consuming and costly to conduct multi-environmental trials and evaluate the CV for new genotypes. Predicting CV based on phenotypes observed in a single environment will greatly reduce the time and cost of the CV evaluation. Although the relationship between plant responses to irrigation changes and drought tolerance has been reported [12,33,34], no study has evaluated the stability of drought tolerance based on phenotypic data collected in a single environmental trial.

This study aimed to investigate a method for evaluating the drought tolerance stability of a genotype based on a single environmental trial. In this study, we set four drought levels, including changing irrigation treatments and conducted 3-year trials. CV was calculated using shoot fresh weights observed in nine combinations of treatments and years. Time-series MS data for each year were modeled using RRMs, and the calculated genetic random regression coefficients were used as secondary traits for the genomic prediction of CV. If genomic prediction models using the calculated genetic random regression coefficients as secondary traits show higher prediction accuracy than those of the simple genomic prediction model without secondary traits, this suggests that time-series MS data are useful in predicting CV. If the genomic prediction models using the time-series MS data collected in the changing irrigation treatments show higher prediction accuracy than those in the continuous drought treatment, this suggests that time-series changes in MS data caused by irrigation change are useful in evaluating drought tolerance stability. Additionally, we built three different prediction models for use in actual breeding schemes: (1) within each combination of treatments and years, (2) using a small dataset of secondary traits, and (3) across years for the same type of treatment. Based on the results of these three cases, we investigated how to predict the CV using the MS data collected in a single environmental trial.

## 2. Materials and Methods

### 2.1. Experimental Data

In this study, the accessions and experimental fields were the same as those used by Sakurai et al. [35]. A diverse panel of 198 soybean accessions were grown in three years 2019, 2020, and 2021 at the Arid Land Research Center, Tottori University, Japan (35°32’ N lat, 134°12’ E long, 14 m above sea level). We used four types of drought treatments: no watering (drought, Treatment D), watering for 5 d followed by no watering for 5 d (Treatment W5), watering for 10 d followed by no watering for 10 d (Treatment W10), and no watering for 10 d followed by watering for 10 d (Treatment D10). Treatments D, W5, and D10 were implemented in 2019. Treatments D, W10, and D10 were implemented in 2020 and 2021 (Figure 1). Each treatment consisted of two rows with one irrigation tube per row, and microplots were placed parallel on either side of the irrigation tubes. For each treatment per year, 198 accessions were randomly assigned to microplots. Each accession was assigned a microplot per treatment. Four plants of each accession were grown in each microplot. The distances between the two rows, microplots, and plants were 50, 80, and 20 cm, respectively. Fertilizers (13, 6.0, 20, 11, 7.0 g/m^2^ of N, P, K, Mg, and Ca, respectively) were applied to the field prior to sowing. Sowing was performed on 10 July 2019, 8 July 2020, and 6 July 2021. Two to three seeds were sown at each position, after which the germinated seedlings were thinned to one per position 2 weeks after sowing.

**Figure 1.**
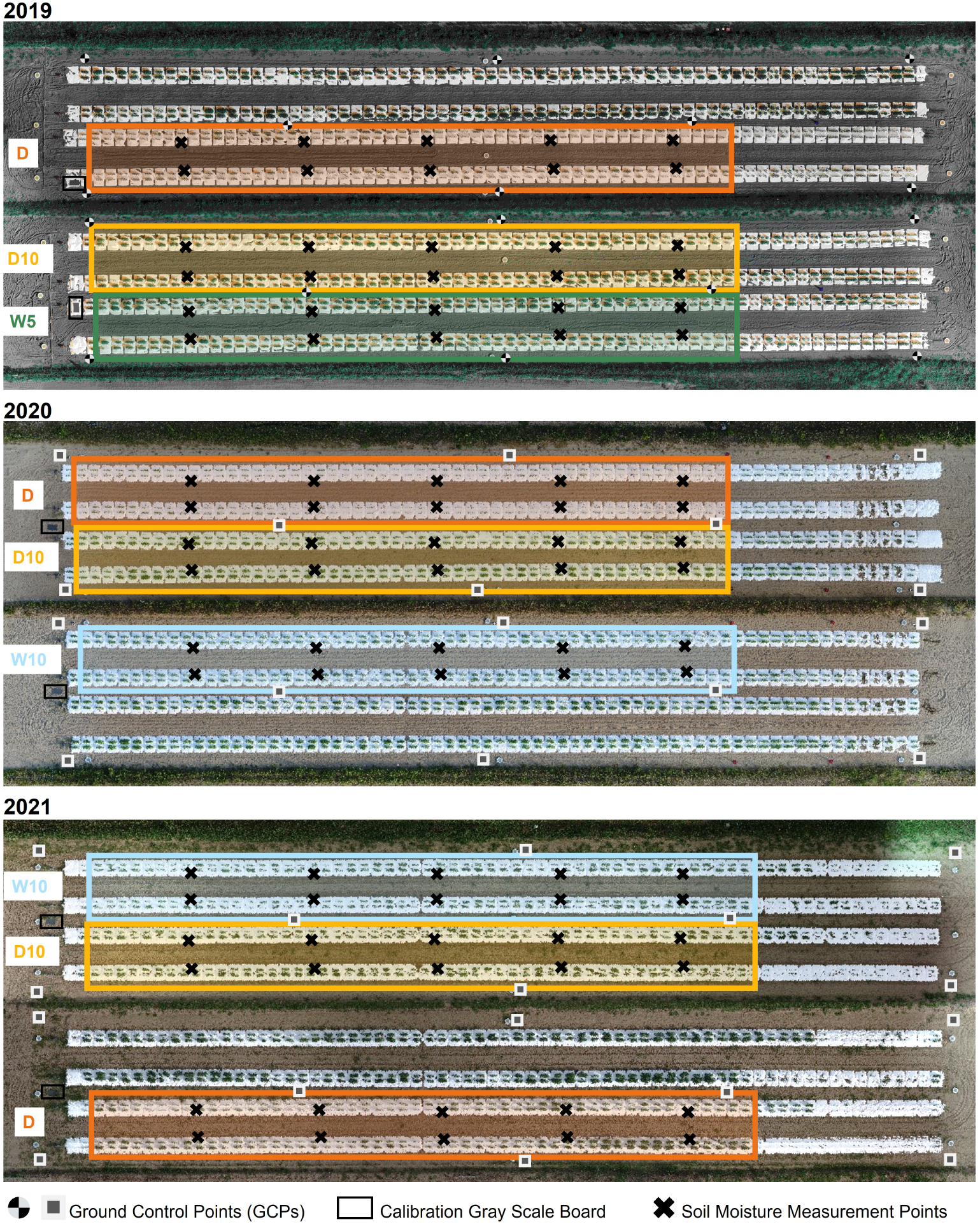
Experimental field overview and treatments in each year. W5: watering for 5 d followed by no watering for 5 d, W10: watering for 10 d followed by no watering for 10 d, D10: no watering for 10 d followed by watering for 10 d, D: no watering treatment.

White mulch sheets (DuPont) were laid over the ridges to prevent rainwater infiltration into the soil and control soil drought levels (Figure 1). A watering tube was installed under the mulch sheets at the center of each row. The watering tube (JKC Agro) irrigated at a flow rate of 1.1 L / h m. Watering was done for over 5 h daily (7:00–9:00, 12:00–14:00, and 16:00–17:00), starting the day after seedling thinning for Treatments W5, W10, and D10. The irrigation cycle for each treatment is shown in Figure S1. Soil moisture was measured using a soil moisture meter (TDR-341F, Fujiwara Seisakusho) at 10 sites for each treatment over 30 d. Except for rainy days, no more than 3 d were allowed between soil moisture measurements. Soil moisture in each treatment was calculated as the average of the treatments in the year (Figure S1). Some accessions did not germinate. Some plants were too small to obtain the MS data, and their measurements were missing. We used 178 accessions with complete data for all the nine treatments (Table S1). In each microplot of the four plants, two centrally located plants were collected 62 d after sowing, and the fresh aboveground weight of each plant was measured. The phenotypic value of the fresh weight of each microplot was calculated as the average of the two plants measured for each microplot.

### 2.2. Multispectral data collection and processing

The MS image collection and image analysis referred to the method employed by Sakurai et al. [35]. In each treatment of a year, MS images were collected using unmanned aerial vehicles (UAVs). In 2019 and 2020, MS images (1.0 cm / pixel) were collected using a four-eye MS camera (Xacti) mounted on a quadcopter UAV (DJI Matrice 100, DJI). The MS camera has four independent lenses and sensors attached to different filters (MidOpt), including a triple-bandpass filter (TB550/660/850) and a red-edge bandpass. The TB550/660/850 can collect spectral intensities at 550 nm (green), 660 nm (red), and 850 nm (near-infrared). Bi725 can collect the spectral intensity at 725 nm (red-edge). The MS camera was set for continuous data capture at two frames per second per lens for a total of eight frames per second. The overlap and sidelap rates were set to 90% and the flights were set to an altitude of 20 m. In 2021, MS images (0.74 cm / pixel) were collected using DJI Phantom 4 Multispectral (P4M). P4M has one RGB sensor and five spectral-band sensors at 450 nm (blue), 560 nm (green), 650 nm (red), 730 nm (red-edge), and 840 nm (near-infrared). P4M continuously collected MS images every two seconds during each flight. The overlap and sidelap rates were set to 75% and the flights were set to an altitude of 15 m.

Each spectral reflectance was calculated as the ratio of each spectral intensity from a grey scale panel set in the experimental field (Figure 1). All the flights were scheduled for 11:00-13:00 under clear sky conditions. Images were collected four, six, and seven times in 2019, 2020, and 2021, respectively (Figure 2). UAVs measurements were scheduled before and after the irrigation treatment was switched on W10 and D10 to capture the response of plants to changes in irrigation. Orthomosaic images of each spectral reflectance were obtained using a Pix4Dmapper (Pix4D). Sixteen ground control points (GCPs) were set up in the field annually. The positions of the GCPs were measured using Aeropoints (Propeller). Using geolocation information from the GCPs, 792 microplots, each with a maximum of four plants, were segmented from the orthomosaic images. Plants were segmented from an image of each microplot using the NDVI-based segmentation method to extract and analyze the MS image data from only the plants. In this experimental field, it was reported that the NDVI differed significantly from the soil surface, white mulch sheets, and plants in 2019 [35]; thus, plants were segmented with the NDVI threshold set to 0.15, as applied by Sakurai et al. [35]. These analyses (extraction of spectral reflectance, calculation of NDVI, and NDVI-based segmentation) were performed using the OpenCV v3.3.1, library in Python v3.6.8.

**Figure 2.**
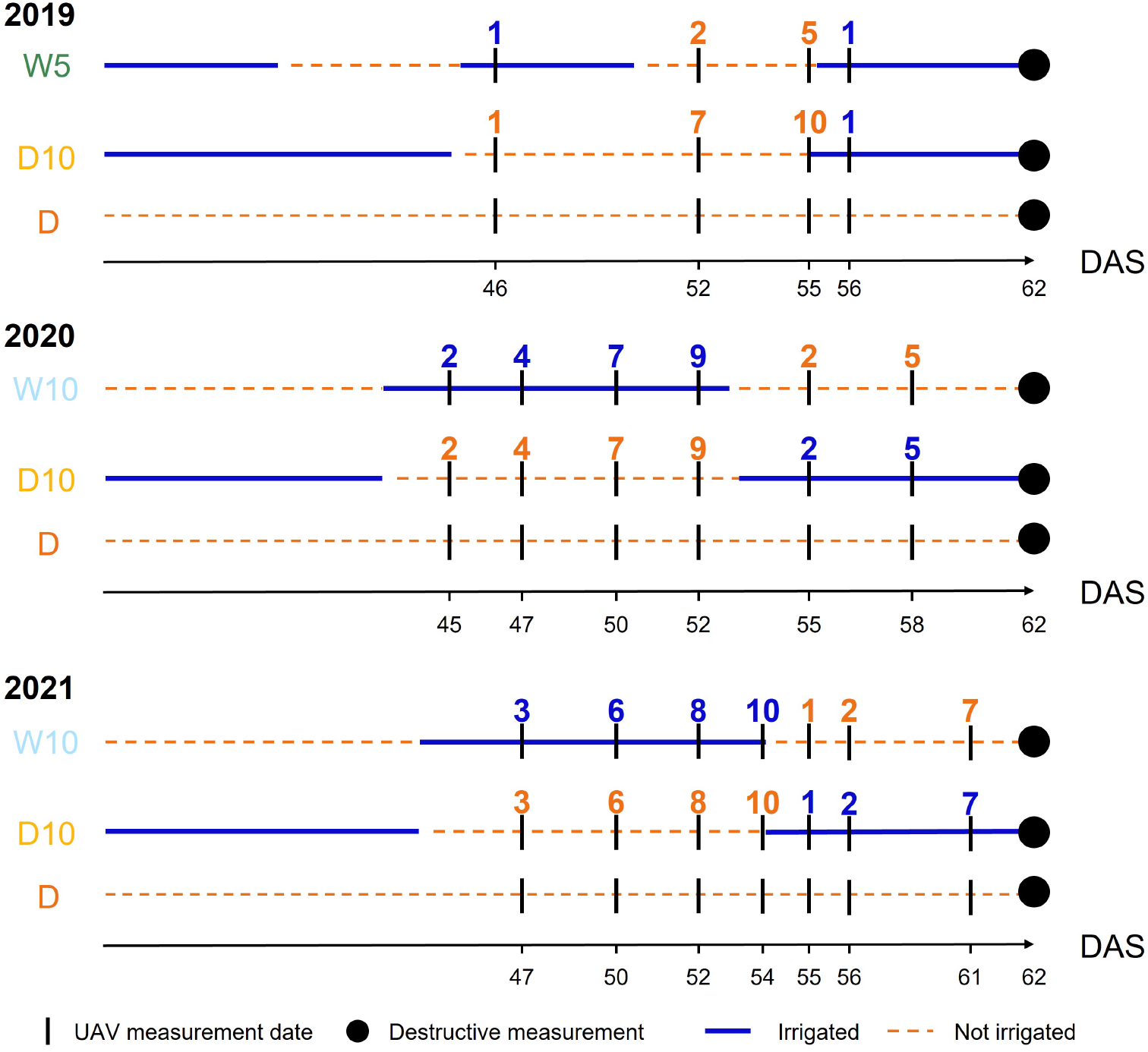
The timings of unmanned aerial vehicle measurements in each year. Blue (respectively Orange) numbers indicate the number of days elapsed since the start of irrigation (respectively drought) treatment. DAS: days after sowing, W5: watering for 5 d followed by no watering 5 d, W10: watering for 10 d followed by no watering 10 d, D10: no watering for 10 d followed by watering 10 d, D: no watering treatment.

Based on the spectral reflectance values of each plant pixel segmented in each microplot, we calculated two types of VIs: NDVI [36,37] and NDRE [38]. The equations for these VIs are listed in Supplementary Table S2. As there were four plants in each microplot, MS data were collected as a community of four plants within each microplot without considering the overlap between plants. Although we attempted to segment only the plant pixels, background noise was not completely removed. In each microplot, the average VI value may have been heavily influenced by the background noise. To reduce this effect, the median of all the segmented plant pixels was used as the representative value of each microplot. The VIs were calculated using R v4.1.2.

### 2.3. Genotyping

The genome dataset was the same as that used by Sakurai et al. [35]. Whole-genome sequence data were obtained for all accessions [39]. All accessions were genotyped using an Illumina HiSeq X Ten or HiSeq 4000 (Illumina), and 4,776,813 single-nucleotide polymorphisms (SNPs) were identified. A total of 173,583 SNP markers were selected from the 4,776,813 SNPs. Using these SNP markers, the additive numerator relationship matrix **G** was estimated using the ‘calcGRM’ function in the ‘RAINBOWR’ package in R v0.1.25 [40].

### 2.4. Coefficient of variation

We defined CV [32] as the environmental stability index for each accession. CV was calculated as follows:

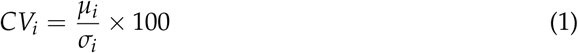

where *μ_i_* is the mean fresh weight in all nine combinations of treatments and years for genotype *i* (*i* = 1,…, 178), *σ_i_* is the standard deviation of the fresh weight in the combinations. A low CV value indicates high environmental stability. Before calculating *μ_i_* and *σ_i_*, fresh weight was scaled by min-max normalization, ranging from 0 to 1 for each combination of treatments and years. This is because *CV_i_* is significantly affected by specific combinations of treatments and years, which have large mean values.

### 2.5. Random regression model

The MS data were collected at four, six, and seven time points in 2019, 2020, and 2021, respectively. A time series of VI values was modeled using a RRM [41] for each treatment of each year separately. The RRM takes the following form:

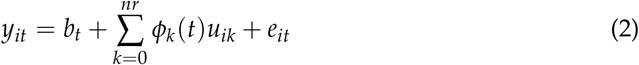

where *y_it_* is the phenotypic value of each VI (NDVI or NDRE) at the time point *t* (*t* = 1,…, 4 in 2019, *t* = 1,…, 6 in 2020, *t* = 1,…, 7 in 2021) for genotype *i* (*i* = 1,…, 178), *b_t_* is the fixed effect of each time point, *nr* is the order of Legendre polynomial for the genetic effects, *ϕ_k_*(*t*) is the *k*th (*k* = 0,…, *nr*) Legendre polynomials for time point *t, u_ik_* is the genetic effect for the *k*th coefficients of Legendre polynomials, and *e_it_* is the random residual effect. Vector 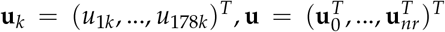 follows the multivariate normal (MVN) distribution, **u** ~ MVN(**0**, **Q** ⊗ **I**_178_) where **Q** is (*nr* + 1) × (*nr* + 1) (co)variance matrix for the Legendre polynomials and **I**_178_ is 178 × 178 identical matrix. Vector 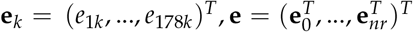 follows the MVN distribution 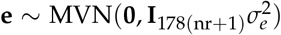, where **I**_178(*nr*+1)_ is the identical matrix and 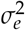 is the residual variance. To determine the order of *nr*, we built RRMs with three values of *nr* (*nr* = 0,1,2) using the data collected in 2021, because the number of time points was the largest in 2021 among other years. The goodness of the model fit was assessed by computing Akaike’s information criterion (AIC) [42]. The best value of *nr* is selected based on the largest AIC value. This RRM model was built using the “ASREML” R package v4.1.0.154 [43].

### 2.6. Genomic heritability and genomic prediction

Simple genomic prediction models were built to calculate the genomic heritability [44] for each trait and to predict the genetic value of the CV. The simple genomic prediction model has the following form:

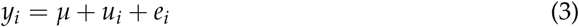

where *y_i_* is the phenotypic value of genotype *i* (*i* = 1,…, 178), *μ* is the overall mean, *u_i_* is the genetic random effect, and *e_i_* is the residual random effect. The vector **u** = (*u*_1_,…, *u*_178_)^*T*^ follows the MVN distribution, 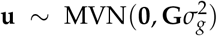 where **G** is the additive numerator relationship matrix, and 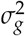 is the additive genetic variance. Vector **e** = (*e*_1_,…,*e*_178_)^*T*^ follows the MVN distribution 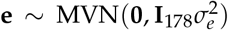, where **I**_178_ is the identical matrix and 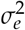 is the residual variance. Based on the estimated parameters of genetic and residual variances, genomic heritability was calculated as 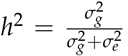. The model was implemented using the ‘EMM.cpp’ function in the ‘RAINBOWR’ package in R v0.1.25 [40].

### 2.7. Multitrait model

For each combination of treatments and years, the MTM was built as a Bayesian multivariate Gaussian model [45,46] to predict the CV of a genotype. An MTM using *M* secondary traits takes the following form:

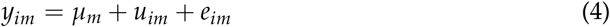

where *y*_*i*0_ is the phenotypic value of the CV for genotype *i* (*i* = 1,…, 178); *y_im_* (*m* = 1,…, *M*) is the phenotypic value of the secondary traits; *μ_m_* is the overall mean for trait *m* (*m* = 0,…, *M*); *u_im_* is the genetic random effect; and *e_im_* is the residual random effect. Vector 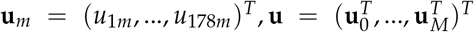 follows the MVN distribution, **u** ~ MVN(**0**, ⊗ ® **G**) where **Σ** is (*M* + 1) × (*M* + 1) genetic (co)variance matrix across traits and **G** is the additive numerator relationship matrix. Vector 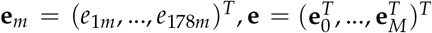 follows the MVN distribution **e** ~ MVN(**0**, **R** ⊗ **I**_178_), where **R** is (*M* + 1) × (*M* + 1) residual (co)variance matrix across traits and **I**_178_ is the identical matrix.

We built two different types of MTMs: (1) an MTM directly using the VI value at each time point as a secondary trait (MT_All_ model), and (2) an MTM using the genetic random regression coefficients of RRM (Equation 2) as a secondary trait (MT_RRM_ model).

### 2.8. Cross-validation cases

We assumed three cases using MTM (Figure 3) and compared the prediction accuracies of MT_All_ model and MT_RRM_ model. The prediction accuracy was evaluated using a 10-fold cross-validation with 10 replicates. Pearson’s correlations were calculated between the observed and predicted CV values in each replicate, and the average of these correlations was used as the prediction accuracy.

**Figure 3.**
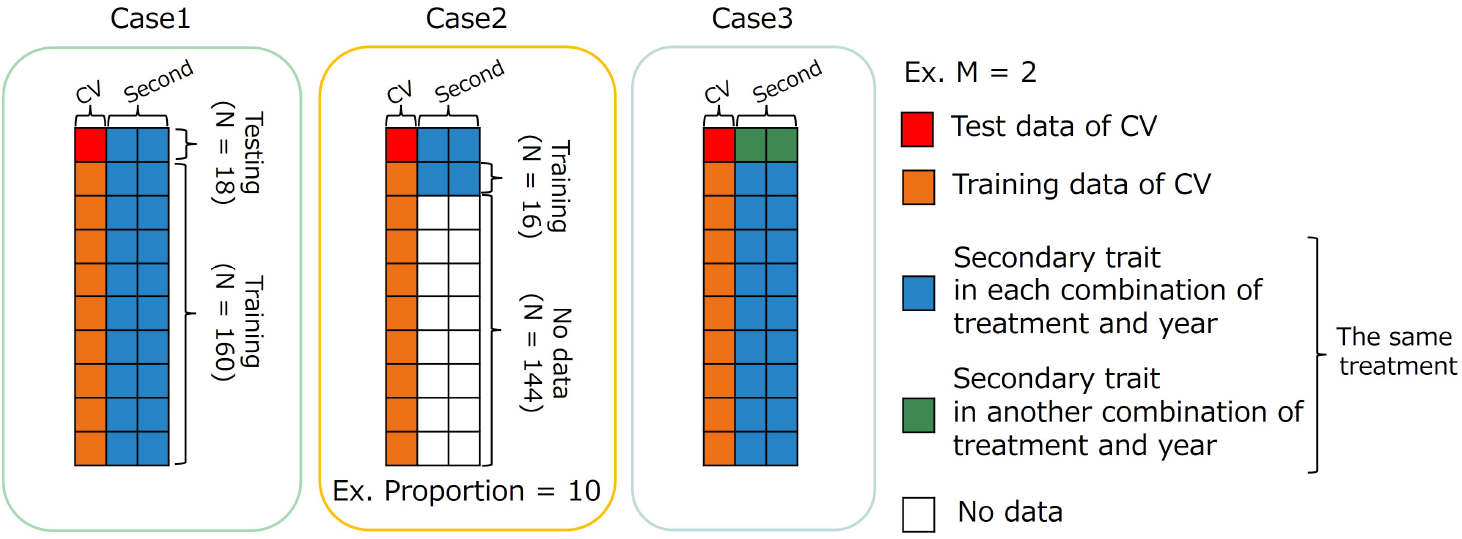
Image representation of three cases using multi-trait (MT) model. The dataset has 178 accessions (N=178). Case1: MT prediction within each combination of treatments and years. Case2: MT prediction within combination of treatments and years with small data set of secondary traits. Case3: MT prediction across years for the same type of treatment. CV: coefficient of variation, M: the number of secondary traits.

#### 2.8.1. Case1: Within each combination of treatments and years

In the first case (Case1), the whole data set in each combination of treatments and years was divided into training and test sets. Predicting the CV of the test set using secondary traits and genome-wide marker data. In MT_All_ model, *M* (the number of secondary traits) was four, six, and seven in 2019, 2020, and 2021, respectively. In MT_RRM_ model, *M* is equal to *nr* + 1 (the order in Equation 2), which is the same for all years.

#### 2.8.2. Case2: Using a small dataset of secondary traits

In the second case (Case2), we assumed that a specific proportion of genotypes had secondary trait data. In MTM, secondary trait data are usually collected for all genotypes in the training and test sets. A small number of genotypes measured for secondary traits in the training set resulted in cost reduction. The proportions of data with secondary traits were set to 10%, 25%, and 50%. Genotypes with secondary trait data were randomly selected five times for each proportion. The prediction accuracy was evaluated via a 10-fold cross-validation with 50 replicates. In MT_All_ model, *M* is four, six, and seven in 2019, 2020, and 2021, respectively. In MT_RRM_ model, *M* is equal to *nr* + 1 for all the years.

#### 2.8.3. Case3: Across years for the same type of treatment

The third case (Case3) was intended to create a prediction model using specific year data and predict CV using data from another year. In this case, we predicted the CV of novel genotypes over the years with their secondary trait data and genome-wide marker data using a previously prepared prediction model. The prediction accuracy was calculated via cross-year cross-validation using the same treatment. Because Treatment W5 did not have yearly replications, this validation was performed only for treatments W10, D10, and D.

In MT_All_ model, *M* is only three because the measurement timings differ among the years. We set the start of irrigation (and drought) in the irrigation changing treatments as a reference point and considered the time difference within a day as the same measurement timing. Therefore, MS data on the dates after sowing (DAS) of 52, 55, and 56 in 2019; 50, 52, and 55 in 2020; and 52, 54, and 55 in 2021 were used in MT_All_ model (Figure 2). However, in MT_RRM_ model, all time point data were available for modeling because time-series MS data were modeled with *nr* + 1 orders, and the genetic random regression coefficients were used as secondary trait data. In MT_RRM_ model, *M* can be fixed at *nr* + 1 for all years.

In all three cases, to compare the prediction accuracy between MT_All_ and MT_RRM_ model we calculated the proportion of improvement. The proportion of improvement is defined as follows:

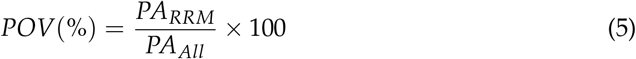

where *POV* denotes the proportion of improvement, *PA_All_* denotes the prediction accuracy of MT_All_ model, and *PA_RRM_* denotes the prediction accuracy of MT_RRM_ model.

## 3. Results

### 3.1. Relationship between coefficient of variation and fresh weight

Fresh weight varied among the nine combinations of treatments and years (Figure 4). As the fresh weight in Treatment D was smaller than that in the other treatments each year, Treatment D was the treatment with the most severe drought stress. For Treatment D10, the fresh weight in 2019 was higher than that in the other two years. In 2019, it rained for 6 out of 10 d from August 25th to September 3rd, the period of no irrigation in Treatment D10. A lack of drought stress during this period might have resulted in the large fresh weight in Treatment D10. Genomic heritability of fresh weight ranged from 0.18 to 0.64 and that of CV was 0.35 (Table 1). Phenotypic correlation between CV and fresh weight in each combination of treatments and years was negative correlation and ranged from −0.45 to −0.12. These results indicate that there is a positive relationship between drought tolerance stability and fresh weight for all combinations of treatments and years.

**Figure 4.**
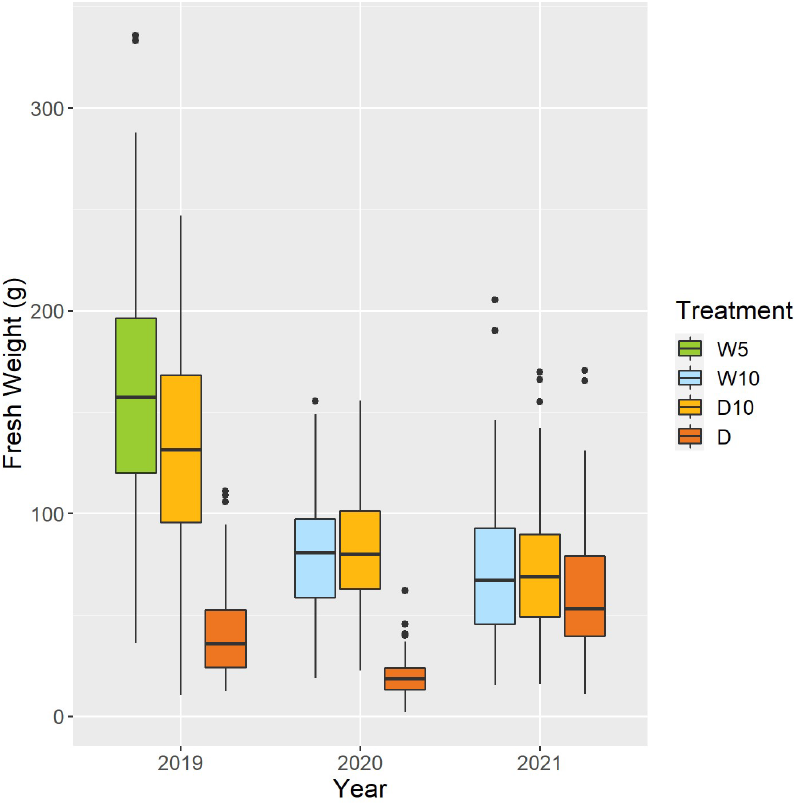
Boxplot of observed fresh weights of 178 soybean accessions in each combination of treatments and years. W5: watering for 5 d followed by no watering 5 d, W10: watering for 10 d followed by no watering 10 d, D10: no watering for 10 d followed by watering 10 d, D: no watering treatment.

**Table 1.** Genomic heritability of fresh weight in each combination of treatments and years, coefficient of variation (CV) calculated over the nine combinations, and phenotypic correlation (r) between fresh weight in each combination of treatments and years and the CV.

| Year | Treatment /Index | $h^2$ | $r$ |
| --- | --- | --- | --- |
| 2019 | W5 | 0.41 | -0.12 |
|  | D10 | 0.23 | -0.12 |
|  | D | 0.23 | -0.35 |
| 2020 | W10 | 0.64 | -0.40 |
|  | D10 | 0.33 | -0.41 |
|  | D | 0.24 | -0.42 |
| 2021 | W10 | 0.18 | -0.44 |
|  | D10 | 0.40 | -0.45 |
|  | D | 0.43 | -0.43 |
|  | CV | 0.35 |  |

\* $h^2$ : genomic heritability, $r$ : phenotypic correlation between fresh weight and coefficient of variation (CV), W5: watering for 5 d followed by no watering 5 d, W10: watering for 10 d followed by no watering 10 d, D10: no watering for 10 d followed by watering 10 d, D: no watering treatment.

### 3.2. Model selection for random regression model

We evaluated the goodness-of-fit of the RRMs using NDVI and NDRE values for each treatment in 2021. Under all treatments, the best model was based on NDVI values using linear Legendre polynomials, that is, the order of *nr* was 1 (Table 2). The model with *nr* = 0 exhibited the best NDRE values under all treatments (Table S3). This result indicates that only the intercept varied among genotypes in the RRM of NDRE values. Therefore, we employed RRM of NDVI using *nr* = 1 in later analyses and calculated the genetic effect for the intercept and 1st coefficients of the Legendre polynomials (L0 and L1) in Equation 2.

**Table 2.** The goodness-of-fit of random regression models (RRMs) with the normalized difference vegetation index (NDVI) values in 2021.

| Treatment | $nr$ | Loglik | AIC | $p$ |
| --- | --- | --- | --- | --- |
| W10 | 0 | 527.2572655 | -1038.514531 | 8 |
|  | **1** | **739.1732966** | **-1458.346593** | **10** |
|  | 2 | 459.0197275 | -892.0394551 | 13 |
| D10 | 0 | 409.6136183 | -803.2272366 | 8 |
|  | **1** | **553.0469996** | **-1086.093999** | **10** |
|  | 2 | 256.6404682 | -487.2809363 | 13 |
| D | 0 | 704.2429918 | -1392.485984 | 8 |
|  | **1** | **725.6443146** | **-1431.288629** | **10** |
|  | 2 | 458.2977954 | -890.5955909 | 13 |

\* $nr$ : the order of Legendre polynomial for the genetic effect, Loglik: log likelihood, AIC: Akaike's information criterion, $p$ : the number of parameters, W10: watering for 10 d followed by no watering 10 d, D10: no watering for 10 d followed by watering 10 d, D: no watering treatment. The best model in each treatment is bolded based on AIC.

### 3.3. Genetic correlation and genomic heritability of secondary traits

We estimated the genetic correlations between CV and L0 and CV and L1, and estimated the genomic heritability of L0, L1, and CV. Except for Treatment W5 in 2019, the genetic correlations between CV and all the traits were negative (Table 3). L0 and L1 values for each genotype were associated with the intercepts and slopes of the time-series NDVI values. Therefore, these negative genetic correlations indicate that a low CV is associated with large intercepts and slopes of the time-series NDVI values. In 2019, there were no significant differences in genetic correlations or genomic heritability among the treatments (Table 3). However, in 2020, treatments W10 and D10 showed higher genetic correlations and genomic heritability than Treatment D for all traits. In 2021, the genetic correlation of L0 was −0.2 in Treatment D, while W10 and D10 showed higher genetic correlations of −0.4 and −0.51, respectively.

**Table 3.**
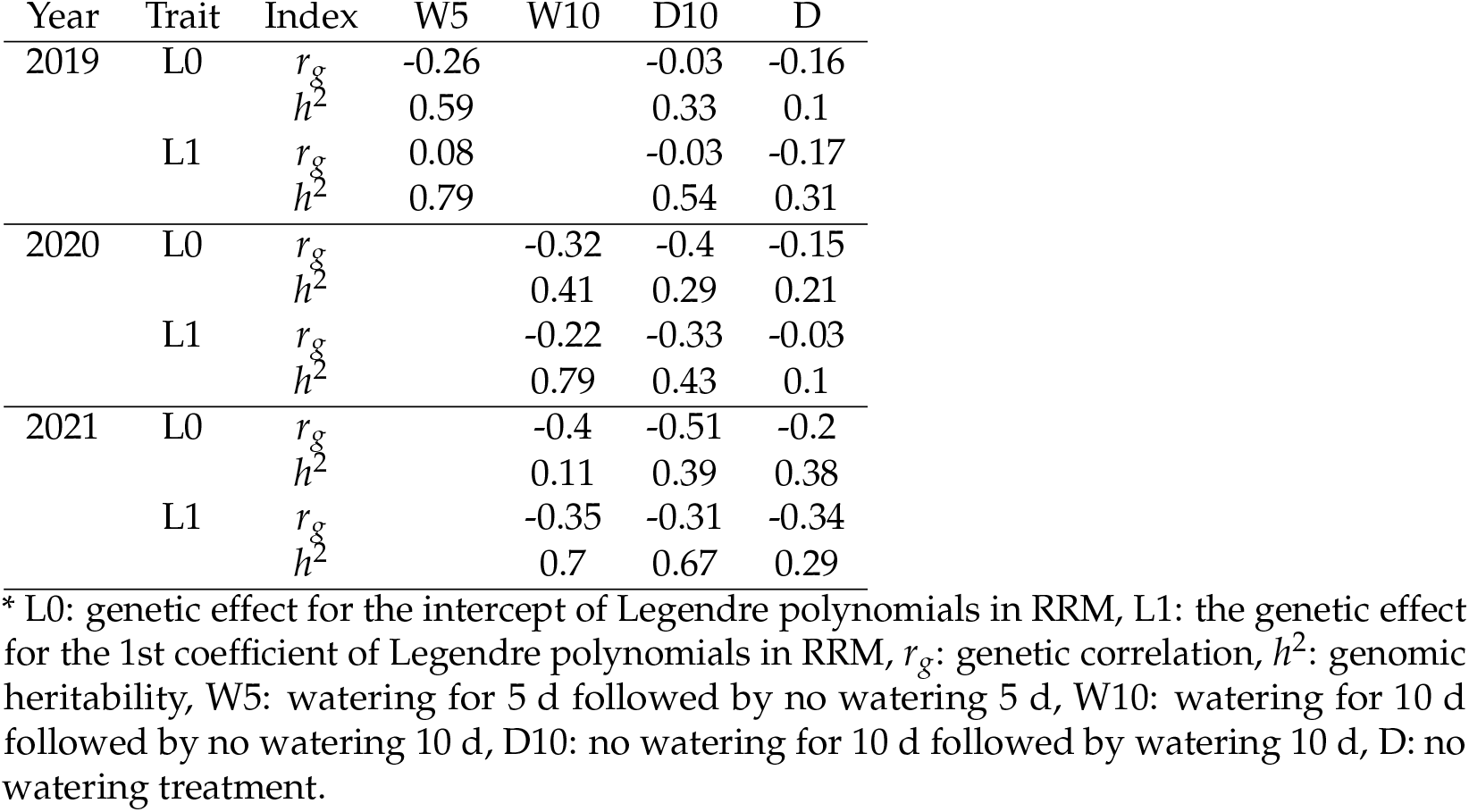
Genetic correlation between each parameter calculated in random regression models (RRMs) and coefficient of variation, and genomic heritability of genetic random regression coefficient of RRMs.

| Year | Trait | Index | W5 | W10 | D10 | D |
| --- | --- | --- | --- | --- | --- | --- |
| 2019 | L0 | $r_g$ | -0.26 | | -0.03 | -0.16 |
| | | $h^2$ | 0.59 | | 0.33 | 0.1 |
| | L1 | $r_g$ | 0.08 | | -0.03 | -0.17 |
| | | $h^2$ | 0.79 | | 0.54 | 0.31 |
| 2020 | L0 | $r_g$ | | -0.32 | -0.4 | -0.15 |
| | | $h^2$ | | 0.41 | 0.29 | 0.21 |
| | L1 | $r_g$ | | -0.22 | -0.33 | -0.03 |
| | | $h^2$ | | 0.79 | 0.43 | 0.1 |
| 2021 | L0 | $r_g$ | | -0.4 | -0.51 | -0.2 |
| | | $h^2$ | | 0.11 | 0.39 | 0.38 |
| | L1 | $r_g$ | | -0.35 | -0.31 | -0.34 |
| | | $h^2$ | | 0.7 | 0.67 | 0.29 |

\* L0: genetic effect for the intercept of Legendre polynomials in RRM, L1: the genetic effect for the 1st coefficient of Legendre polynomials in RRM, $r_g$ : genetic correlation, $h^2$ : genomic heritability, W5: watering for 5 d followed by no watering 5 d, W10: watering for 10 d followed by no watering 10 d, D10: no watering for 10 d followed by watering 10 d, D: no watering treatment.

### 3.4. Case1: Within each combination of treatments and years

In Case1, the prediction accuracy of MT_RRM_ model was higher than that of the simple genomic prediction model for all year treatments (Figure 5a). In 2019, there was no difference in the prediction accuracy among the treatments. However, the prediction accuracies of treatments W10 and D10 in 2020 were 25% and 30% higher, respectively than those of Treatment D in 2020. In addition, the prediction accuracies of treatments W10 and D10 in 2021 were higher by 14% and 11%, respectively than that of Treatment D in 2021. These results indicate that the time-series MS data collected during the treatment, which changed the irrigation pattern, were more useful than those in the continuous drought treatment for predicting the CV in 2020 and 2021. To evaluate the importance of using RRMs, we compared MT_RRM_ model and MT_All_ model. In the seven combinations of treatments and years, the prediction accuracies of MT_RRM_ model were higher than those of MT_All_ model (Figure 5b). In particular, the proportion of improvement is 14% for Treatment W10 in 2021.

**Figure 5.**
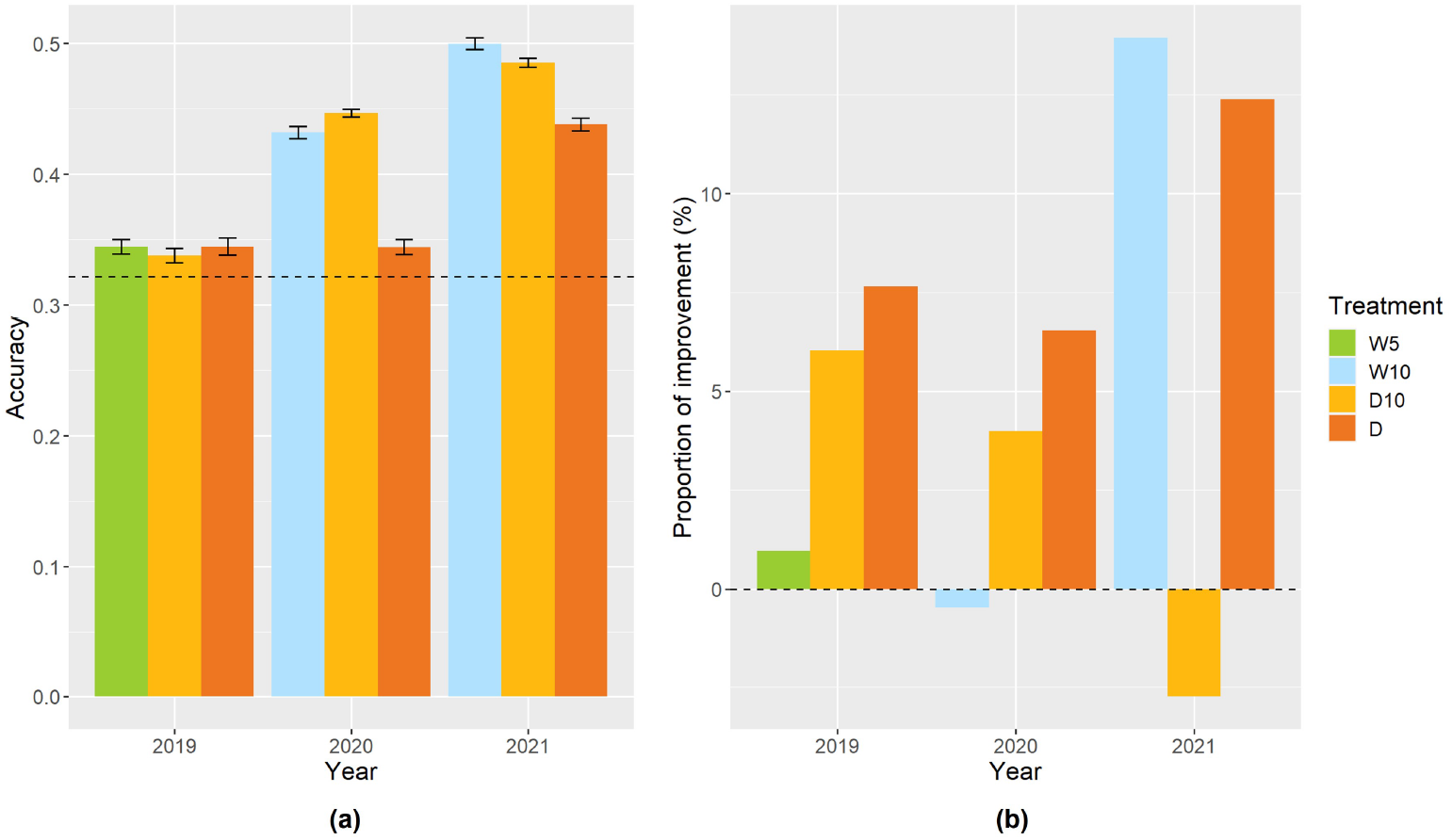
Prediction accuracy of MT_RRM_ model and the comparison of MT_RRM_ model and MT_All_ model in Case1. Error bars represent standard error over 10 replicate cross-validations. (**a**) Prediction accuracy of MT_RRM_ model within each combination of treatments and years. A dashed line represents the prediction accuracy of the simple genomic prediction model. (**b**) The proportion of improvement calculated using the prediction accuracy of MT_RRM_ model and MT_All_ model. W5: watering for 5 d followed by no watering 5 d, W10: watering for 10 d followed by no watering 10 d, D10: no watering for 10 d followed by watering 10 d, D: no watering treatment.

### 3.5. Case2: Using a small dataset of secondary traits

To reduce the cost and labor required to create the prediction model, we changed the proportion of genotypes collected as secondary trait data in the training set. In 2020 and 2021, except for Treatment D in 2020, the prediction accuracy for all combinations of treatments and years increased as the proportion of genotypes with secondary trait data increased (Figure 6). In Treatment D10 of 2020 and 2021, even when the proportion was 10 percent, the prediction accuracies of MT_RRM_ model were 23% and 30% higher, respectively, than those of the simple genomic prediction model. The proportion of improvement was close to zero, indicating that there was no significant difference between MT_RRM_ model and MT_All_ model (Figure S2).

**Figure 6.**
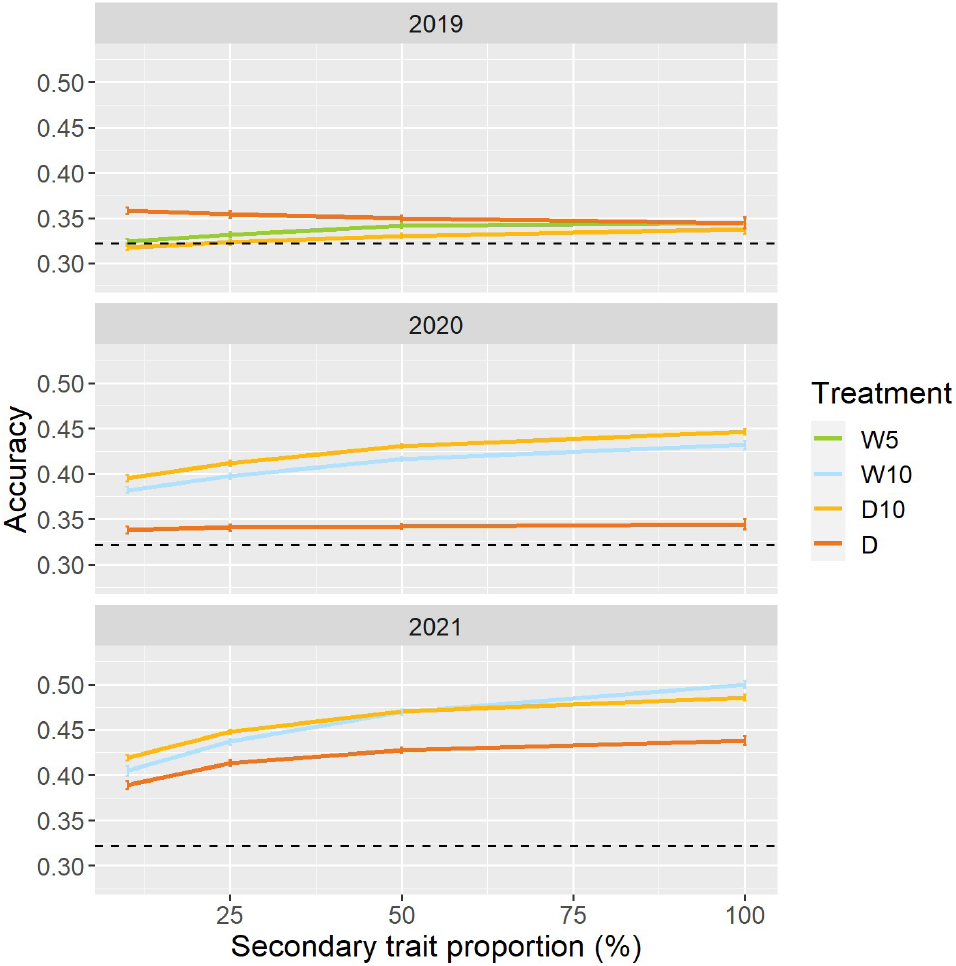
Prediction accuracy of MT_RRM_ model when a specific proportion of genotypes in the training set did not have secondary trait data. Dashed line represents the prediction accuracy of simple genomic prediction model. Error bars represent standard error over 50 replicate crossvalidations. W5: watering for 5 d followed by no watering 5 d, W10: watering for 10 d followed by no watering 10 d, D10: no watering for 10 d followed by watering 10 d, D: no watering treatment.

### 3.6. Case3: Across years for the same type of treatment

In Case3, we evaluated the across-year predictions for each treatment. In all three treatments, MT_RRM_ models outperformed the simple genomic prediction model (Figure 7a). In cross-validations with the 2020 and 2021 datasets, the prediction accuracy of MT_RRM_ models for Treatments W10 and D10 were on average 42% and 42% higher than that of the simple genomic prediction model, respectively. The proportion of improvement in all MT_RRM_ models was greater than zero (Figure 7b). The proportion of improvement was, on average, 11, 6, and 6% for treatments W10, D10, and D, respectively.

**Figure 7.**
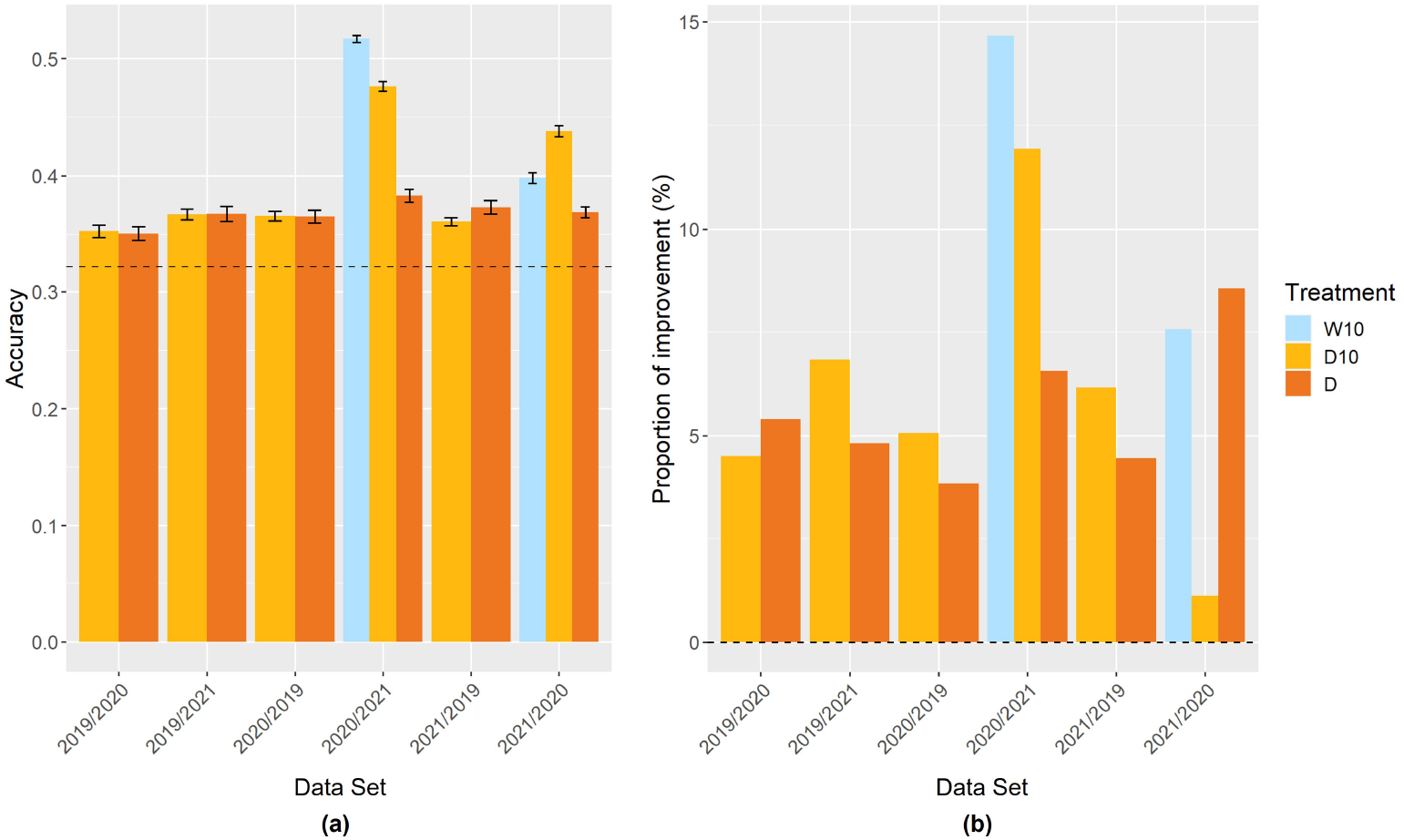
Prediction accuracy of MT_RRM_ models and the proportion of improvement in Case3. Item names, such as 2019/2020, represent the training year and test year. (**a**) Prediction accuracy of MT_RRM_ model among years in the same treatment. Dashed line represents the prediction accuracy of simple genomic prediction model. Error bars represent standard error over 10 replicate crossvalidations. (**b**) The proportion of improvement in each combination. W5: watering for 5 d followed by no watering 5 d, W10: watering for 10 d followed by no watering 10 d, D10: no watering for 10 d followed by watering 10 d, D: no watering treatment.

## 4. Discussion

### 4.1. Usefulness of MS data for predicting drought tolerance stability

In this study, we compared the prediction accuracy of a simple genomic prediction model and a genomic prediction model using time-series NDVI values as secondary traits to predict the CV (Figure 5,6,7). In soybeans, a relationship between NDVI and drought tolerance has been reported [7]. Except for Case2 in 2019, all genomic prediction models using time-series NDVI values as a secondary trait showed higher prediction accuracy than a simple genomic prediction model (Figure 5,6,7). These results suggest that time-series MS data are useful for predicting the CV.

In 2020 and 2021, the prediction accuracy of treatments W10 and D10, which changed irrigation treatments, was higher than that of Treatment D, which was the no watering treatment, in the same year (Figure 5,6). A relationship between plant responses to changes in irrigation and drought tolerance has been reported. In soybeans, slow wilting resulted in drought tolerance [2,3,47,48] and a relationship between slow wilting and NDVI values under drought is observed [7]. Additionally, the speed of recovery from drought stress is associated with drought tolerance in several crop species [49,50], including soybean [51]. However, no study has evaluated the drought tolerance stability using time-series NDVI changes caused by changes in irrigation. This result indicates that plant responses to irrigation changes are useful for evaluating drought tolerance stability.

In this study, the overall prediction accuracy of CV was lower in 2019 than that in the other years. One possible reason for this is that NDVI was measured in 2019 less frequently than in 2020 and 2021. NDVI was measured six and seven times in 2020 and 2021, respectively; however, in 2019, NDVI was measured only four times. The small number of measurements may not capture the time-series changes in NDVI well. In addition, we collected NDVI values on the first day after the irrigation change in Treatment D10 in 2019 (Figure 1). Therefore, it is difficult to capture plant responses to changes in irrigation.

### 4.2. The advantage of using random regression

In Case1, MT_RRM_ model, which modeled time-series NDVI data and used L0 and L1 (genetic random regression coefficients) as secondary traits in the MTM, generally showed a higher prediction accuracy than MT_All_ model, which treated time-series NDVI data as independent traits and used each day’s NDVI value as a secondary trait in the MTM (Figure 5). It was reported that the RRM is superior to the MTM in modeling time-series data [52,53]. In addition, there are problems with over-parameterization and high computational requirements when treating time-series data as independent traits and constructing MTMs [11,54,55]. In terms of calculation efficiency, RRM is useful for modeling time-series data.

Campbell et al. [13] built RRM using time-series projected shoot area (PSA) data for 179 rice lines obtained in their own study. Using the built-in RRM [13], Campbell predicted time-series changes in PSA for new 178 rice lines collected in a different year [56]. This result suggests that RRM is useful for predictions across years. In our study, MT_RRM_ model is superior to MT_All_ model in terms of prediction across years in Case3 (Figure 7). MT_RRM_ model can build a model using all-day data of NDVI value in each year, whereas MT_All_ model can only use dates that are common across all years. This difference in the amount of data may be the reason for the difference in prediction accuracy. When using MT_All_ model for predictions across years, it is necessary to match the data measurement dates. It was difficult to collect data on a desired date because of weather conditions or equipment problems. At this point, the RRM can be built without considering the measurement date and number of days of data measurement.

### 4.3. Utility for breeding

Environmental stability, that is, the stability of phenotypes over environments, is an important target in breeding [57–60], and the CV of phenotypic values over environments can be a good index for selecting stable genotypes. To evaluate the environmental stability, it is necessary to conduct multi-environmental experiments which require high costs and labor [61–64]. In this study, we propose a scheme for evaluating environmental stability using only one environmental experiment. The results of Case1 validation suggest that drought tolerance stability can be evaluated in a single environment by capturing plant responses to irrigation changes (Figure 5). The results of Case2 validation indicate that even when 10% of genotypes in a training set had secondary trait data, MT_RRM_ model was more accurate than a simple genomic prediction model in treatments W10 and D10 in 2020 and 2021 (Figure 6). Therefore, it is possible to reduce the number of genotypes measured for secondary traits in the training set. The results of Case3 validation also suggest that we can predict stability across years (Figure 7). Once a prediction model is constructed, the stability of novel genotypes can be predicted based on time-series NDVI data. The results of Case2 and Case3 validations also demonstrate the flexibility of MT_RRM_ model. When the drought tolerance stability of novel genotypes is evaluated in a single environment, it greatly reduces the time and cost, and thus streamlines breeding schemes.

In this study, we attempted to build a scheme for modeling the relationships among phenotypes, plant responses to irrigation changes, and drought tolerance stability. HTP can capture the time-series plant responses to environmental and irrigation changes [12,33]. The analysis of plant responses to environmental changes is expected to provide important insights that have not yet been obtained [11,65]. This study provides new insights that the drought tolerance stability of each genotype is related to time-series NDVI changes in each genotype caused by the irrigation changes.

## Supplementary Materials

The following supporting information can be downloaded at: https://www.mdpi.com/article/10.3390/1010000/s1, Figure S1: The cycle of irrigation treatment and the day change of soil moisture content in each combination of treatments and years; Figure S2: Comparison of prediction accuracy between MT_RRM_ model and MT_RRM_ model in Case2. Table S1: Description of the 178 accessions used in this study; Table S2: Equations for the two vegetation indices (VIs) used in this study; Table S3: The goodness-of-fit of random regression models (RRMs) with the normalized difference red-edge (NDRE) values in 2021.

## Author Contributions

Methodology, software, formal analysis, data curation, writing—original draft preparation and visualization, K.S.; investigation, data curation and project administration, Y.T.; methodology and software, K.M.; conceptualization, investigation, data curation, writing—review and editing, supervision, project administration and funding acquisition, H.I.; Conceptualization, resources, data curation, and funding acquisition, A.K.; Conceptualization, data curation, and funding acquisition, Y.O., Y.Y., H.T.(Hirokazu Takahashi), H.T.(Hideki Takanashi), M.T., H.T.(Hisashi Tsujimoto), M.N. and T.F. All authors have read and agreed to the published version of the manuscript.

## Institutional Review Board Statement

Not applicable.

## Informed Consent Statement

Not applicable.

## Data Availability Statement

The datasets generated and analyzed in the present study are available from the ‘Sakuraikengo/RRMTM_supple’ repository in the GitHub, https://github.com/Sakuraikengo/RRMTM_supple.

## Acknowledgments

We are grateful to the technical staff of Arid Land Research Center, Tottori University, and Izumi Higashida.

## Abbreviations

The following abbreviations are used in this manuscript:

CV: Coefficient of variation
RRM: Random regression model
HTP: High throughput phenotyping
HS: Hyperspectral
MS: Multispectral
NDVI: Normalized difference vegetation index
NDRE: Normalized difference red-edge
VIs: Vegetation indices
GWAS: Genome wide association study
MTM: Multi-trait model
UAV: Unmanned aerial vehicle
P4M: Phantom 4 Multispectral
SNP: Single-nucleotide polymorphism
MVN: Multivariate normal
AIC: Akaike’s information criterion

## Disclaimer/Publisher’s Note

The statements, opinions and data contained in all publications are solely those of the individual author(s) and contributor(s) and not of MDPI and/or the editor(s). MDPI and/or the editor(s) disclaim responsibility for any injury to people or property resulting from any ideas, methods, instructions or products referred to in the content.

## References

1. Specht, J.E.; Hume, D.J.; Kumudini, S.V. Soybean yield potential - A genetic and physiological perspective. Crop Science. 1999, 39, 1560–1570. https://doi.org/10.2135/cropsci1999.3961560x.

2. Pathan, S.M.; Lee, J.D.; Sleper, D.A.; Fritschi, F.B.; Sharp, R.E.; Carter, T.E.; Nelson, R.L.; King, C.A.; Schapaugh, W.T.; Ellersieck, M.R.; et al. Two soybean plant introductions display slow leaf wilting and reduced yield loss under drought. Journal of Agronomy and Crop Science. 2014, 200, 231–236. https://doi.org/10.1111/jac.12053.

3. Ye, H.; Song, L.; Schapaugh, W.T.; Ali, M.L.; Sinclair, T.R.; Riar, M.K.; Mutava, R.N.; Li, Y.; Vuong, T.; Valliyodan, B.; et al. The importance of slow canopy wilting in drought tolerance in soybean. Journal of Experimental Botany. 2020, 71, 642–652. https://doi.org/10.1093/jxb/erz150.

4. Bi, Y.; Kong, W.; Huang, W. Hyperspectral diagnosis of nitrogen status in arbuscular mycorrhizal inoculated soybean leaves under three drought conditions. International Journal of Agricultural and Biological Engineering. 2018, 11, 126–131. https://doi.org/10.25165/j.ijabe.20181106.4019.

5. Wijewardana, C.; Alsajri, F.A.; Irby, J.T.; Krutz, L.J.; Golden, B.; Henry, W.B.; Gao, W.; Reddy, K.R. Physiological assessment of water deficit in soybean using midday leaf water potential and spectral features. Journal of Plant Interactions. 2019, 14, 533–543. https://doi.org/10.1080/17429145.2019.1662499.

6. Winterhalter, L.; Mistele, B.; Jampatong, S.; Schmidhalter, U. High throughput phenotyping of canopy water mass and canopy temperature in well-watered and drought stressed tropical maize hybrids in the vegetative stage. European Journal of Agronomy. 2011, 35, 22–32. https://doi.org/10.1016/j.eja.2011.03.004.

7. Zhou, J.; Zhou, J.; Ye, H.; Ali, M.L.; Nguyen, H.T.; Chen, P. Classification of soybean leaf wilting due to drought stress using UAV-based imagery. Computers and Electronics in Agriculture. 2020, 175, 105576. https://doi.org/10.1016/j.compag.2020.105576.

8. Du, W.; Yu, D.; Fu, S. Detection of quantitative trait loci for yield and drought tolerance traits in soybean using a recombinant inbred line population. Journal of Integrative Plant Biology. 2009, 51, 868–878. https://doi.org/10.1111/j.1744-7909.2009.00855.x.

9. Zygielbaum, A.I.; Gitelson, A.A.; Arkebauer, T.J.; Rundquist, D.C. Non-destructive detection of water stress and estimation of relative water content in maize. Geophysical Research Letters. 2009, 36, 2–5. https://doi.org/10.1029/2009GL038906.

10. Yang, G.; Liu, J.; Zhao, C.; Li, Z.; Huang, Y.; Yu, H.; Xu, B.; Yang, X.; Zhu, D.; Zhang, X.; et al. Unmanned aerial vehicle remote sensing for field-based crop phenotyping: Current status and perspectives. Frontiers in Plant Science. 2017, 8, 1111. https://doi.org/10.3389/fpls.2017.01111.

11. Moreira, F.F.; Oliveira, H.R.; Volenec, J.J.; Rainey, K.M.; Brito, L.F. Integrating High-Throughput Phenotyping and Statistical Genomic Methods to Genetically Improve Longitudinal Traits in Crops. Frontiers in Plant Science. 2020, 11, 1–18. https://doi.org/10.3389/fpls.2020.00681.

12. Chen, D.; Neumann, K.; Friedel, S.; Kilian, B.; Chen, M.; Altmann, T.; Klukas, C. Dissecting the phenotypic components of crop plant growthand drought responses based on high-throughput image analysis. Plant Cell. 2014, 26, 4636–4655. https://doi.org/10.1105/tpc.114.129601.

13. Campbell, M.; Walia, H.; Morota, G. Utilizing random regression models for genomic prediction of a longitudinal trait derived from high-throughput phenotyping. Plant Direct. 2018, 2, 1–11. https://doi.org/10.1002/pld3.80.

14. Oliveira, H.R.; Brito, L.F.; Lourenco, D.A.; Silva, F.F.; Jamrozik, J.; Schaeffer, L.R.; Schenkel, F.S. Invited review: Advances and applications of random regression models: From quantitative genetics to genomics. Journal of Dairy Science. 2019, 102, 7664–7683. https://doi.org/10.3168/jds.2019-16265.

15. Freitas Moreira, F.; Rojas de Oliveira, H.; Lopez, M.A.; Abughali, B.J.; Gomes, G.; Cherkauer, K.A.; Brito, L.F.; Rainey, K.M. High-Throughput Phenotyping and Random Regression Models Reveal Temporal Genetic Control of Soybean Biomass Production. Frontiers in Plant Science. 2021, 12, 715983. https://doi.org/10.3389/fpls.2021.715983.

16. Henderson, C.R. Applications of linear models in animal breeding; University of Guelph: Guelph, 1984.

17. Huisman, A.E.; Veerkamp, R.F.; Van Arendonk, J.A. Genetic parameters for various random regression models to describe the weight data of pigs. Journal of Animal Science. 2002, 80, 575–582. https://doi.org/10.2527/2002.803575x.

18. Schaeffer, L.R. Application of random regression models in animal breeding. Livestock Production Science. 2004, 86, 35–45. https://doi.org/10.1016/S0301-6226(03)00151-9.

19. Campbell, M.; Momen, M.; Walia, H.; Morota, G. Leveraging Breeding Values Obtained from Random Regression Models for Genetic Inference of Longitudinal Traits. The Plant Genome. 2019, 12, 180075. https://doi.org/10.3835/plantgenome2018.10.0075.

20. Sun, J.; Rutkoski, J.E.; Poland, J.A.; Crossa, J.; Jannink, J.L.; Sorrells, M.E. Multitrait, Random Regression, or Simple Repeatability Model in High-Throughput Phenotyping Data Improve Genomic Prediction for Wheat Grain Yield. The Plant Genome. 2017, 10, plantgenome2016.11.0111. https://doi.org/10.3835/plantgenome2016.11.0111.

21. Kumar, A.; Verulkar, S.B.; Mandal, N.P.; Variar, M.; Shukla, V.D.; Dwivedi, J.L.; Singh, B.N.; Singh, O.N.; Swain, P.; Mall, A.K.; et al. High-yielding, drought-tolerant, stable rice genotypes for the shallow rainfed lowland drought-prone ecosystem. Field Crops Research. 2012, 133, 37–47. https://doi.org/10.1016/j.fcr.2012.03.007.

22. Pidgeon, J.D.; Ober, E.S.; Qi, A.; Clark, C.J.; Royal, A.; Jaggard, K.W. Using multi-environment sugar beet variety trials to screen for drought tolerance. Field Crops Research. 2006, 95, 268–279. https://doi.org/10.1016/j.fcr.2005.04.010.

23. Torres, R.O.; Henry, A. Yield stability of selected rice breeding lines and donors across conditions of mild to moderately severe drought stress. Field Crops Research. 2018, 220, 37–45. https://doi.org/10.1016/j.fcr.2016.09.011.

24. Kumar, S.; Kumari, J.; Bansal, R.; Kuri, B.R.; Upadhyay, D.; Srivastava, A.; Rana, B.; Yadav, M.K.; Sengar, R.S.; Singh, A.K.; et al. Multi-environmental evaluation of wheat genotypes for drought tolerance. Indian Journal of Genetics and Plant Breeding. 2018, 78, 26–35. https://doi.org/10.5958/0975-6906.2018.00004.4.

25. Ayed, S.; Othmani, A.; Bouhaouel, I.; da Silva, J.A. Multi-environment screening of durum wheat genotypes for drought tolerance in changing climatic events. Agronomy. 2021, 11, 1–15. https://doi.org/10.3390/agronomy11050875.

26. Das, S.; Misra, R.C.; Patnaik, M.C.; Das, S.R. GxE interaction, adaptability and yield stability of mid-early rice genotypes. Indian Journal of Agricultural Research. 2010, 44, 104–111.

27. Di Matteo, J.A.; Ferreyra, J.M.; Cerrudo, A.A.; Echarte, L.; Andrade, F.H. Yield potential and yield stability of Argentine maize hybrids over 45 years of breeding. Field Crops Research. 2016, 197, 107–116. https://doi.org/10.1016/j.fcr.2016.07.023.

28. Mohammadi, R.; Amri, A. Comparison of parametric and non-parametric methods for selecting stable and adapted durum wheat genotypes in variable environments. Euphytica. 2008, 159, 419–432. https://doi.org/10.1007/s10681-007-9600-6.

29. Rao, M.R.; Willey, R.W. Evaluation of yield stability in intercropping: Studies on sorghum/pigeonpea. Experimental Agriculture. 1980, 16, 105–116. https://doi.org/10.1017/S0014479700010796.

30. Küchenmeister, F.; Küchenmeister, K.; Wrage, N.; Kayser, M.; Isselstein, J. Yield and yield stability in mixtures of productive grassland species: Does species number or functional group composition matter? Grassland Science. 2012, 58, 94–100. https://doi.org/10.1111/j.1744-697X.2012.00242.x.

31. Ray, D.K.; Gerber, J.S.; Macdonald, G.K.; West, P.C. Climate variation explains a third of global crop yield variability. Nature Communications. 2015, 6, 1–9. https://doi.org/10.1038/ncomms6989.

32. Francis.; Kannenberg. Yield stability studies in short-season maize. I. A descriptive method for grouping genotypes. Canadian Journal of plant science. 1978, 58, 1029–1034. https://doi.org/10.4141/cjps78-157.

33. Dodig, D.; Božinović, S.; Nikolić, A.; Zorić, M.; Vančetović, J.; Ignjatović-Micić, D.; Delić, N.; Weigelt-Fischer, K.; Altmann, T.; Junker, A. Dynamics of Maize Vegetative Growth and Drought Adaptability Using Image-Based Phenotyping Under Controlled Conditions. Frontiers in Plant Science. 2021, 12, 1–18. https://doi.org/10.3389/fpls.2021.652116.

34. Marchetti, C.F.; Ugena, L.; Humplík, J.F.; Polák, M.; Ćavar Zeljković, S.; Podlešáková, K.; Fürst, T.; De Diego, N.; Spíchal, L. A Novel Image-Based Screening Method to Study Water-Deficit Response and Recovery of Barley Populations Using Canopy Dynamics Phenotyping and Simple Metabolite Profiling. Frontiers in Plant Science. 2019, 10, 1–20. https://doi.org/10.3389/fpls.2019.01252.

35. Sakurai, K.; Toda, Y.; Kajiya-Kanegae, H.; Ohmori, Y.; Yamasaki, Y.; Takahashi, H.; Takanashi, H.; Tsuda, M.; Tsujimoto, H.; Kaga, A.; et al. Time-series multispectral imaging in soybean for improving biomass and genomic prediction accuracy. Plant Genome. 2022, 15, e20244. https://doi.org/10.1002/tpg2.20244.

36. Price, J.C.; Bausch, W.C. Leaf area index estimation from visible and near-infrared reflectance data. Remote Sensing of Environment. 1995, 52, 55–65. https://doi.org/10.1016/0034-4257(94)00111-Y.

37. Zarate-Valdez, J.L.; Metcalf, S.; Stewart, W.; Ustin, S.L.; Lampinen, B. Potentials and limits of vegetation indices for LAI and APAR assessment. Precision agriculture. 2015, 16, 161–173. https://doi.org/10.2136/sssaj1977.03615995004100040037x.

38. Jorge, J.; Vallbé, M.; Soler, J.A. Detection of irrigation inhomogeneities in an olive grove using the NDRE vegetation index obtained from UAV images. European Journal of Remote Sensing. 2019, 52, 169–177. https://doi.org/10.1080/22797254.2019.1572459.

39. Kajiya-Kanegae, H.; Nagasaki, H.; Kaga, A.; Hirano, K.; Ogiso-Tanaka, E.; Matsuoka, M.; Ishimori, M.; Ishimoto, M.; Hashiguchi, M.; Tanaka, H.; et al. Whole-genome sequence diversity and association analysis of 198 soybean accessions in mini-core collections. DNA Research. 2021, 28, 1–13. https://doi.org/10.1093/dnares/dsaa032.

40. Hamazaki, K.; Iwata, H. Rainbow: Haplotype-based genome-wide association study using a novel SNP-set method. PLoS Computational Biology. 2020, 16, 1–17. https://doi.org/10.1371/journal.pcbi.1007663.

41. Mrode, R.A. Linear Models For The Prediction Of Animal Breeding Values; Cabi, 2014. https://doi.org/doi.org/10.1079/9781780643915.0000.

42. Hirotugu Akaike. A New Look at the Statistical Model Identification; Springer Series in Statistics: New York, 1974. https://doi.org/10.1007/978-1-4612-1694-0_16.

43. Gilmour, a.R.; Gogel, B.J.; Cullis, B.R.; Welham, S.J.; Thompson, R. ASReml User Guide Release 4.1 Structural Specification; Hemel hempstead:, 2015.

44. Litchfield, K.; Thomsen, H.; Mitchell, J.S.; Sundquist, J.; Houlston, R.S.; Hemminki, K.; Turnbull, C. Quantifying the heritability of testicular germ cell tumour using both population-based and genomic approaches. Scientific Reports. 2015, 5, 1–7. https://doi.org/10.1038/srep13889.

45. Lado, B.; Vázquez, D.; Quincke, M.; Silva, P.; Aguilar, I.; Gutiérrez, L. Resource allocation optimization with multi-trait genomic prediction for bread wheat (Triticum aestivum L.) baking quality. Theoretical and Applied Genetics. 2018, 131, 2719–2731. https://doi.org/10.1007/s00122-018-3186-3.

46. Montesinos-López, O.A.; Montesinos-López, A.; Crossa, J.; Toledo, F.H.; Pérez-Hernández, O.; Eskridge, K.M.; Rutkoski, J. A genomic bayesian multi-trait and multi-environment model. G3: Genes, Genomes, Genetics. 2016, 6, 2725–2774. https://doi.org/10.1534/g3.116.032359.

47. Fletcher, A.L.; Sinclair, T.R.; Allen, L.H. Transpiration responses to vapor pressure deficit in well watered ‘slow-wilting’ and commercial soybean. Environmental and Experimental Botany. 2007, 61, 145–151. https://doi.org/10.1016/j.envexpbot.2007.05.004.

48. Valliyodan, B.; Ye, H.; Song, L.; Murphy, M.; Grover Shannon, J.; Nguyen, H.T. Genetic diversity and genomic strategies for improving drought and waterlogging tolerance in soybeans. Journal of Experimental Botany. 2017, 68, 1835–1849. https://doi.org/10.1093/jxb/erw433.

49. Hayano-Kanashiro, C.; Calderón-Vásquez, C.; Ibarra-Laclette, E.; Herrera-Estrella, L.; Simpson, J. Analysis of gene expression and physiological responses in three Mexican maize landraces under drought stress and recovery irrigation. PLoS ONE. 2009, 4, e7531. https://doi.org/10.1371/journal.pone.0007531.

50. Kränzlein, M.; Geilfus, C.M.; Franzisky, B.L.; Zhang, X.; Wimmer, M.A.; Zörb, C. Physiological Responses of Contrasting Maize (Zea mays L.) Hybrids to Repeated Drought. Journal of Plant Growth Regulation. 2021, 41, 2708–2718. https://doi.org/10.1007/s00344-021-10468-2.

51. Hossain, M.M.; Liu, X.; Qi, X.; Lam, H.M.; Zhang, J. Differences between soybean genotypes in physiological response to sequential soil drying and rewetting. Crop Journal. 2014, 2, 366–380. https://doi.org/10.1016/j.cj.2014.08.001.

52. Kranis, A.; Su, G.; Sorensen, D.; Woolliams, J.A. The application of random regression models in the genetic analysis of monthly egg production in turkeys and a comparison with alternative longitudinal models. Poultry Science. 2007, 86, 470–475. https://doi.org/10.1093/ps/86.3.470.

53. Mota, R.R.; Marques, L.F.; Lopes, P.S.; da Silva, L.P.; Neto, F.R.; de Resende, M.D.; Torres, R.A. Genetic evaluation using multi-trait and random regression models in Simmental beef cattle. Genetics and Molecular Research. 2013, 12, 2465–2480. https://doi.org/10.4238/2013.July.24.2.

54. Misztal, I.; Strabel, T.; Jamrozik, J.; Mäntysaari, E.A.; Meuwissen, T.H. Strategies for estimating the parameters needed for different test-day models. Journal of Dairy Science. 2000, 83, 1125–1134. https://doi.org/10.3168/jds.S0022-0302(00)74978-2.

55. Oh, S.H.; See, M.T. Comparison of genetic parameter estimates of total sperm cells of boars between random regression and multiple trait animal models. Asian-Australasian Journal of Animal Sciences. 2008, 21, 923–927. https://doi.org/10.5713/ajas.2008.70383.

56. Campbell, M.T.; Du, Q.; Liu, K.; Brien, C.J.; Berger, B.; Zhang, C.; Walia, H. A Comprehensive Image-based Phenomic Analysis Reveals the Complex Genetic Architecture of Shoot Growth Dynamics in Rice (Oryza sativa). The Plant Genome. 2017, 10,plantgenome2016.07.0064. https://doi.org/10.3835/plantgenome2016.07.0064.

57. Blum, A. Plant breeding for stress environments; CRC press, 2018; p. 231. https://doi.org/10.1201/9781351075718.

58. Sabaghnia, N.; Sabaghpour, S.H.; Dehghani, H. The use of an AMMI model and its parameters to analyse yield stability in multi-environment trials. Journal of Agricultural Science. 2008, 146, 571–581. https://doi.org/10.1017/S0021859608007831.

59. Mickelbart, M.V.; Hasegawa, P.M.; Bailey-Serres, J. Genetic mechanisms of abiotic stress tolerance that translate to crop yield stability. Nature Reviews Genetics. 2015, 16, 237–251. https://doi.org/10.1038/nrg3901.

60. Tollenaar, M.; Lee, E.A. Yield potential, yield stability and stress tolerance in maize. Field Crops Research. 2002, 75, 161–169. https://doi.org/10.1016/S0378-4290(02)00024-2.

61. Fan, X.M.; Kang, M.S.; Chen, H.; Zhang, Y.; Tan, J.; Xu, C. Yield stability of maize hybrids evaluated in multi-environment trials in Yunnan, China. Agronomy Journal. 2007, 99, 220–228. https://doi.org/10.2134/agronj2006.0144.

62. Karimizadeh, R.; Mohammadi, M.; Sabaghni, N.; Mahmoodi, A.A.; Roustami, B.; Seyyedi, F.; Akbari, F. GGE Biplot Analysis of Yield Stability in Multi-environment Trials of Lentil Genotypes under Rainfed Condition. Notulae Scientia Biologicae. 2013, 5, 256–262. https://doi.org/10.15835/nsb529067.

63. Lal, R.K.; Chanotiya, C.S.; Gupta, P.; Mishra, A. Influences of traits associations for essential oil yield stability in multi-environment trials of vetiver (Chrysopogon zizanioides L. Roberty). Biochemical Systematics and Ecology. 2022, 103, 104448. https://doi.org/10.1016/j.bse.2022.104448.

64. Akter, A.; Hasan, M.J.; Kulsum, U.; Rahman, M.; Khatun, M.; Islam, M. GGE Biplot Analysis for Yield Stability in Multi-environment Trials of Promising Hybrid Rice (Oryza sativa L.). Bangladesh Rice Journal. 2015, 19, 1–8. https://doi.org/10.3329/brj.v19i1.25213.

65. Arnold, P.A.; Kruuk, L.E.; Nicotra, A.B. How to analyse plant phenotypic plasticity in response to a changing climate. New Phytologist. 2019, 222, 1235–1241. https://doi.org/10.1111/nph.15656.

