## Supplementary for "Evaluating drought tolerance stability in soybean by the response of irrigation change captured from time-series multispectral data"

(a)

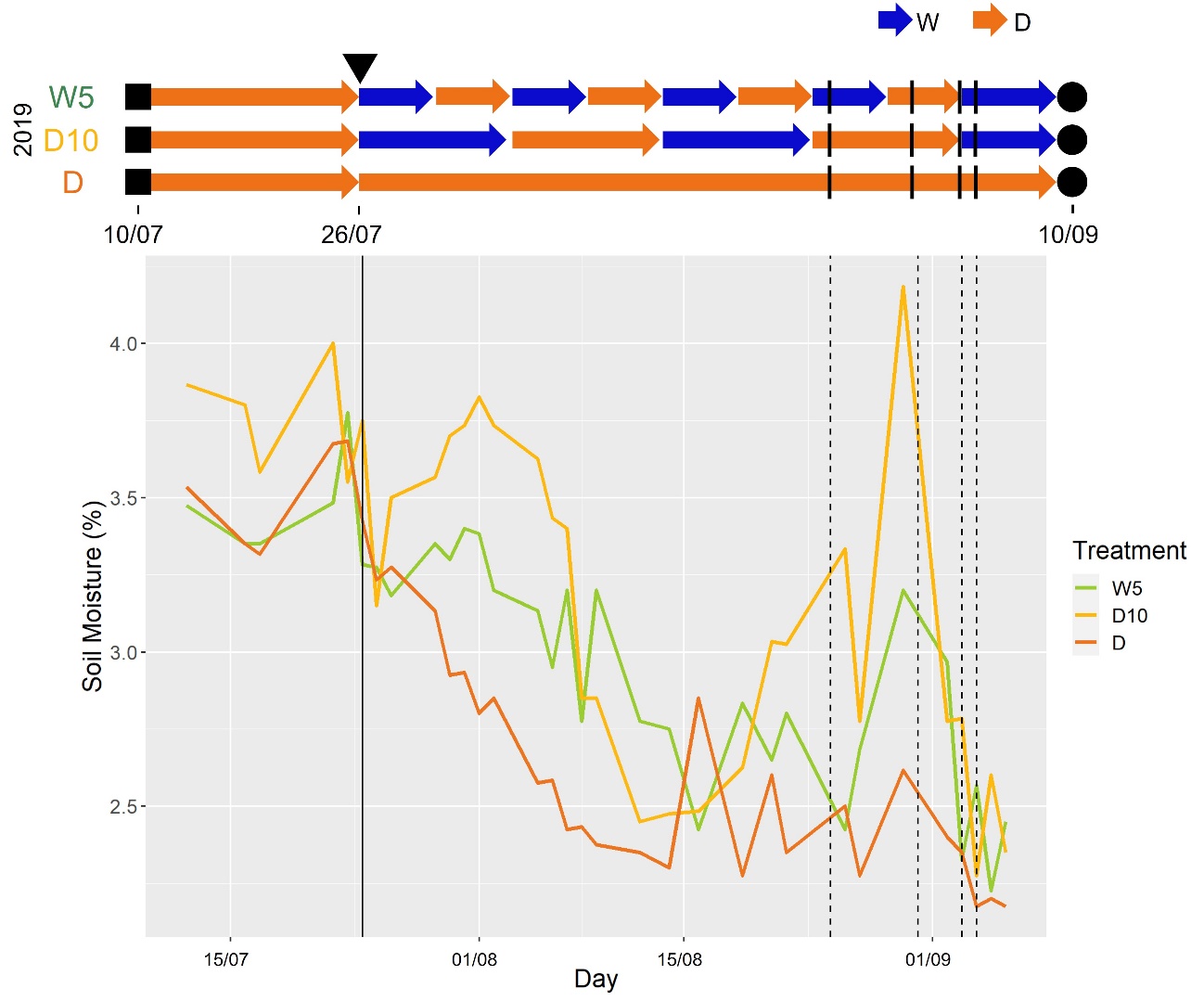

(b)

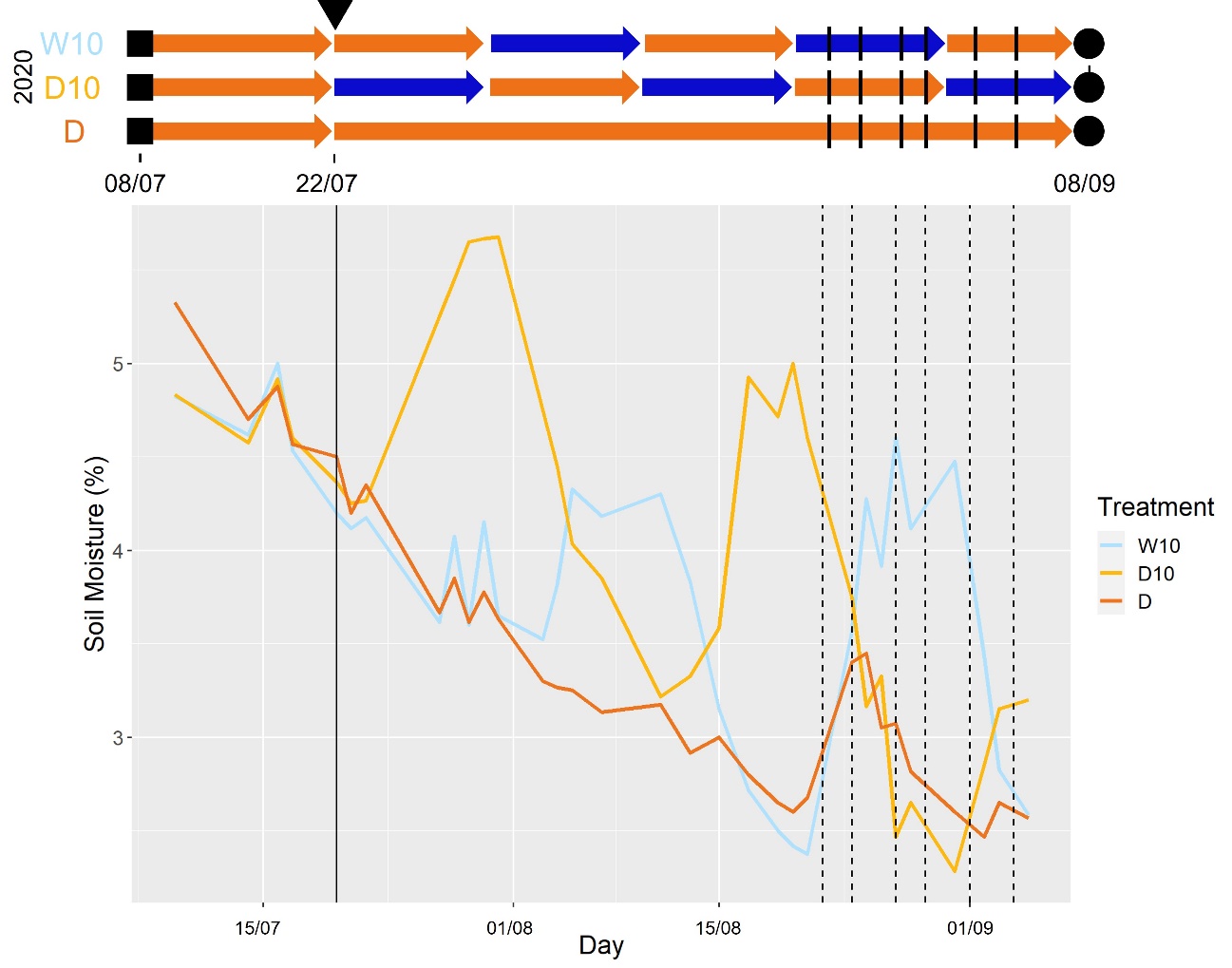

(c)

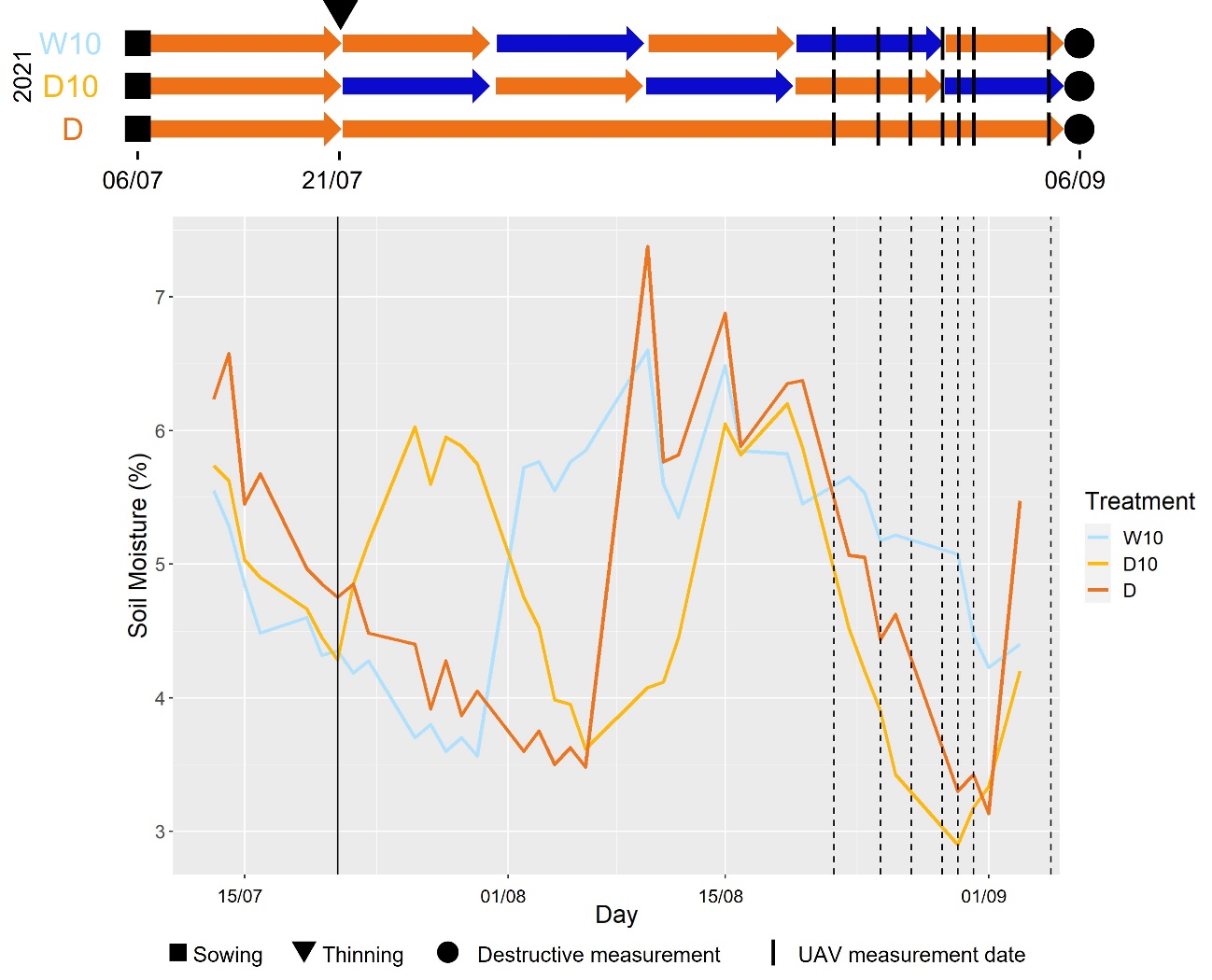

Figure S1.	The cycle of irrigation treatment and the day change of soil moisture content in each combination of treatments and years. W5: watering for 5 d followed by no watering 5 d, W10: watering for 10 d followed by no watering 10 d, D10: no watering for 10 d followed by watering 10 d, D: no watering treatment. Two colors of arrows mean the irrigation treatment: not irrigated period (orange), irrigated period (blue). The solid line represents the date of thinning and dashed lines represent the date of UAV measurements. (a) in 2019, (b) in 2020, (c) in 2021.

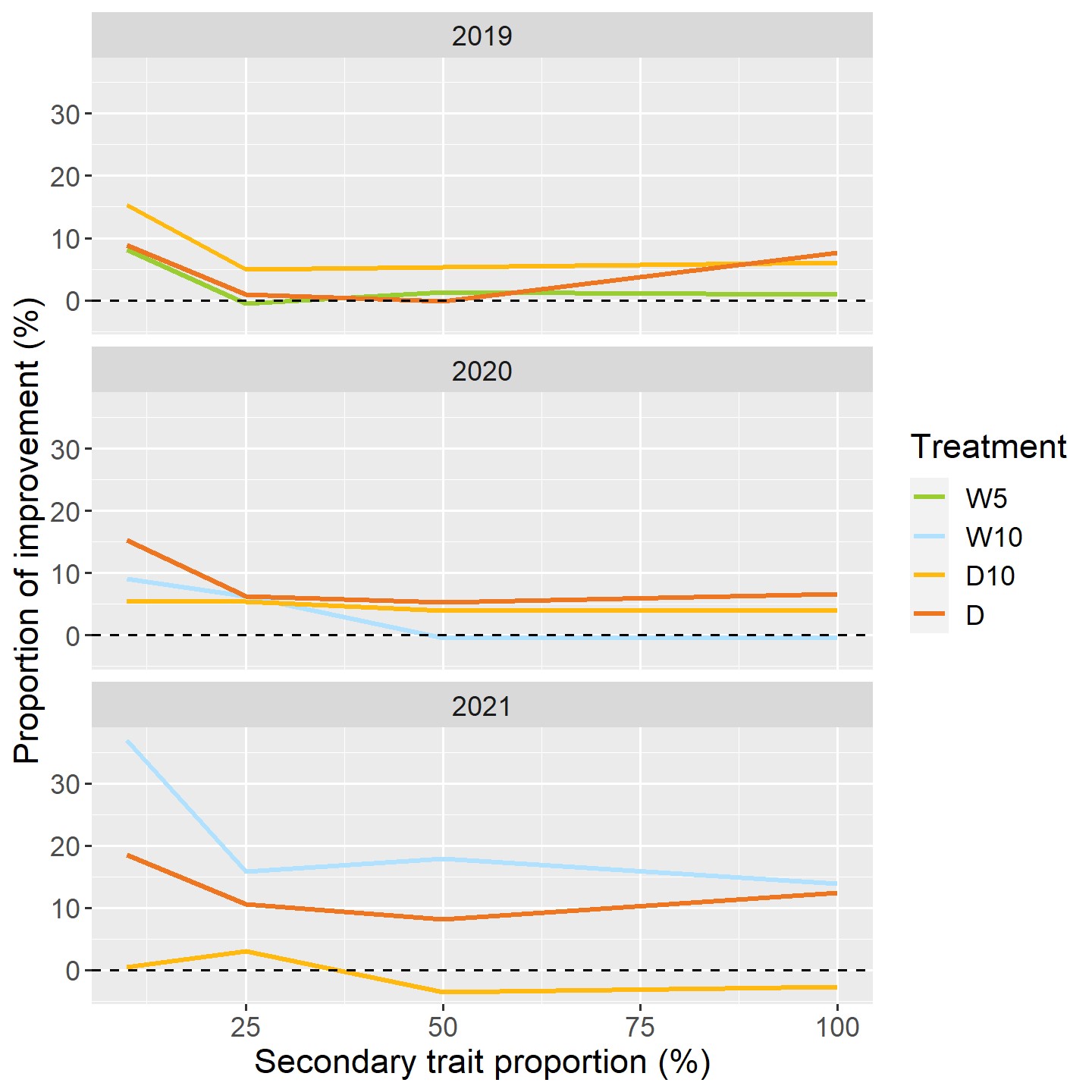

Figure S2.	The comparison of prediction accuracy between $\mathrm{MT}_{\mathrm{RRM}}$ model and $\mathrm{MT}_{\mathrm{All}}$ model in Case2. W5: watering for 5 d followed by no watering 5 d, W10: watering for 10 d followed by no watering 10 d, D10: no watering for 10 d followed by watering 10 d, D: no watering treatment.

**Table S1	The description of 178 accessions that are used in this study. We used soybean genetic resources, registered as the mini core collections in the National Institute of Agrobiological Sciences (NIAS) gene bank. Some accessions were removed due to the unavailability of phenotypic traits.**

| 5002T | C1329 | GmJMC003 | GmJMC004 | GmJMC007 |
| --- | --- | --- | --- | --- |
| GmJMC009 | GmJMC013 | GmJMC016 | GmJMC017 | GmJMC021 |
| GmJMC023 | GmJMC025 | GmJMC026 | GmJMC028 | GmJMC030 |
| GmJMC031 | GmJMC032 | GmJMC033 | GmJMC034 | GmJMC037 |
| GmJMC039 | GmJMC040 | GmJMC041 | GmJMC044 | GmJMC047 |
| GmJMC049 | GmJMC050 | GmJMC051 | GmJMC053 | GmJMC054 |
| GmJMC055 | GmJMC056 | GmJMC057 | GmJMC058 | GmJMC059 |
| GmJMC060 | GmJMC061 | GmJMC062 | GmJMC063 | GmJMC064 |
| GmJMC065 | GmJMC067 | GmJMC068 | GmJMC069 | GmJMC076 |
| GmJMC077 | GmJMC078 | GmJMC079 | GmJMC080 | GmJMC082 |
| GmJMC085 | GmJMC091 | GmJMC092 | GmJMC093 | GmJMC095 |
| GmJMC096 | GmJMC097 | GmJMC098 | GmJMC099 | GmJMC100 |
| GmJMC101 | GmJMC102 | GmJMC104 | GmJMC105 | GmJMC106 |
| GmJMC110 | GmJMC111 | GmJMC112 | GmJMC114 | GmJMC116 |
| GmJMC117 | GmJMC121 | GmJMC126 | GmJMC128 | GmJMC130 |
| GmJMC131 | GmJMC133 | GmJMC137 | GmJMC139 | GmJMC145 |
| GmJMC149 | GmJMC158 | GmJMC161 | GmJMC167 | GmJMC172 |
| GmJMC177 | GmJMC179 | GmJMC180 | GmJMC184 | GmWMC001 |
| GmWMC006 | GmWMC010 | GmWMC011 | GmWMC012 | GmWMC014 |
| GmWMC015 | GmWMC018 | GmWMC022 | GmWMC024 | GmWMC027 |
| GmWMC029 | GmWMC035 | GmWMC036 | GmWMC038 | GmWMC042 |
| GmWMC045 | GmWMC046 | GmWMC048 | GmWMC066 | GmWMC070 |
| GmWMC071 | GmWMC072 | GmWMC073 | GmWMC074 | GmWMC075 |
| GmWMC083 | GmWMC084 | GmWMC086 | GmWMC089 | GmWMC094 |
| GmWMC103 | GmWMC107 | GmWMC108 | GmWMC109 | GmWMC115 |
| GmWMC118 | GmWMC119 | GmWMC120 | GmWMC122 | GmWMC123 |
| GmWMC124 | GmWMC125 | GmWMC127 | GmWMC129 | GmWMC132 |
| GmWMC134 | GmWMC135 | GmWMC136 | GmWMC140 | GmWMC141 |
| GmWMC142 | GmWMC143 | GmWMC144 | GmWMC146 | GmWMC147 |
| GmWMC148 | GmWMC151 | GmWMC152 | GmWMC153 | GmWMC154 |
| GmWMC155 | GmWMC156 | GmWMC159 | GmWMC160 | GmWMC162 |
| GmWMC163 | GmWMC164 | GmWMC165 | GmWMC166 | GmWMC168 |
| GmWMC169 | GmWMC171 | GmWMC173 | GmWMC174 | GmWMC175 |
| GmWMC176 | GmWMC178 | GmWMC181 | GmWMC182 | GmWMC183 |
| GmWMC185 | GmWMC186 | GmWMC188 | GmWMC189 | GmWMC190 |
| GmWMC191 | GmWMC192 | Houjaku Kuwazu |  |  |

**Table S2**	**The equations of two vegetation indices (VIs) which are used in this study.**

| **Index** | **Equation** |
| --- | --- |
| Normalized difference vegetation index (NDVI) | $(\rho NIR-\rho RED)/$  $(\rho NIR+\rho RED)$ |
| Normalized difference red-edge index (NDRE) | $(\rho NIR-\rho RE)/$  $(\rho NIR+\rho RE)$ |

Note: $\rho RED, \rho RE, \rho NIR$ represent the spectral reflectance of red (660 nm or 650 nm), red-edge (725 nm or 730 nm), and near-infrared (850 nm or 840 nm).

**Table S3**	**The goodness-of-fit of random regression models (RRMs) with the normalized difference red-edge (NDRE) values in 2021.**

| Treatment | $nr$ | Loglik | AIC | $p$ |
| --- | --- | --- | --- | --- |
| **W10** | **0** | **794.7129** | **-1573.43** | **8** |
| W10 | 1 | 731.8879 | -1443.78 | 10 |
| W10 | 2 | 425.5259 | -825.052 | 13 |
| **D10** | **0** | **598.1608** | **-1180.32** | **8** |
| D10 | 1 | 554.6491 | -1089.3 | 10 |
| D10 | 2 | 308.1588 | -590.318 | 13 |
| **D** | **0** | **696.3551** | **-1376.71** | **8** |
| D | 1 | 539.5232 | -1059.05 | 10 |
| D | 2 | 437.6705 | -849.341 | 13 |

* $nr$: the order of Legendre polynomial for the genetic effect, Loglik: log likelihood, AIC: Akaike’s information criterion, $p$: the number of parameters, W10: watering for 10 d followed by no watering 10 d, D10: no watering for 10 d followed by watering 10 d, D: no watering treatment. The best model in each treatment is bolded.
